# Electrobehavioral phenotype and seizure pharmacosensitivity in a novel mouse model of patient-derived *SLC6A1* S295L mutation-associated neurodevelopmental epilepsy

**DOI:** 10.1101/2021.12.17.473036

**Authors:** Britta E. Lindquist, Yuliya Voskobiynyk, Jeanne T. Paz

## Abstract

Solute carrier family 6 member 1 (*SLC6A1*) gene encodes GAT-1, a GABA transporter expressed on glia and presynaptic terminals of inhibitory neurons. Mutations in *SLC6A1* are associated with myoclonic atonic epilepsy, absence epilepsy, autism, and intellectual disability. However, the mechanisms leading to these defects are unknown. Here, we used a novel mouse model harboring a point mutation (S295L) recently identified in the human *SLC6A1* gene that results in impaired membrane trafficking of the GAT-1 protein. We performed chronic wireless telemetry recordings of heterozygous (GAT-1^S295L/+^) mice, and of mice lacking one or both copies of the gene (GAT-1^+/−^ and GAT-1^−/−^). We assessed their behaviors and pharmacosensitivity, and investigated the relationship between seizure burden and behavioral performance. GAT-1^S295L/+^ mice exhibited frequent spike-wave discharges (SWDs) associated with behavioral arrest, and there was a dose-effect relationship between GAT-1 gene copy number and the severity of electrocorticogram (ECoG) abnormalities. Seizure burden was inversely correlated with behavioral performance. Forelimb grip strength was reduced in female mice. Acute administration of GAT-1 antagonist NO-711 induced SWDs in wildtype mice, exacerbated the phenotype in GAT-1^S295L/+^ and GAT-1^+/−^ mice, and had no effect on GAT-1^−/−^ mice lacking the drug target. By contrast, ethosuximide normalized the ECoG in GAT-1^S295L/+^ and GAT-1^+/−^ mice. In conclusion, GAT-1^S295L/+^ mice show haploinsufficiency with evidence of GAT-1 hypofunction. This mouse model reconstitutes major aspects of human disease and thus provides a useful preclinical model for drug screening and gene therapy.

**Significance Statement:** The *SLC6A1* gene encodes for GAT-1, a major GABA transporter. Mutations in *SLC6A1* lead to a spectrum of neurodevelopmental disorders and epilepsies collectively referred to as *SLC6A1*-related disorders (SRD). A critical contributor to disability in SRD patients is the burden of seizures and their sequelae. There is an urgent need to understand the mechanisms of SRD as they will inform therapeutic interventions. Mouse models can provide critical information allowing both the assessment of candidate therapies and the design of next generations therapies. Here we used behavioral assessments and wireless electrophysiology in a new mouse model of SRD to understand the disease pathogenesis and the association between seizure burden and behavioral deficits.

## INTRODUCTION

The solute carrier family 6 member 1 (*SLC6A1*) gene encodes the GABA transporter 1 (GAT-1), a sodium- and chloride-dependent transporter that regulates extracellular GABA levels. GAT-1 is expressed mainly in astrocytes and the terminals of GABAergic neurons, where it regulates GABA levels in the synaptic and extrasynaptic compartments (Conti et al., 1998; Kersanté et al., 2013). Patients with mutations in the *SLC6A1* gene have myoclonic atonic and absence seizures, sleep disturbances, attention deficit hyperactivity disorder, motor deficits, developmental delay, and autism, collectively referred to as *SLC6A1*-related disorders (SRD) (Carvill et al., 2015; Johannesen et al., 2018; Wang et al., 2020; Goodspeed et al., 2020; Kahen et al., 2021; Anon, 2021). The prognosis is poor (Johannesen et al., 2018; Goodspeed et al., 2020). In a recent compilation of clinical cases (Goodspeed et al., 2020), the most commonly recorded clinical features are epilepsy (in 91% of cases) and developmental delay (in 82% of cases). Seizure onset is early (mean 2.5 years) and exacerbates developmental delay (Lal, 2021). The incidence of SRD is estimated to be 2.65 per 100,000 births, although this number is expected to increase once *SLC6A1* is included in standard clinical epilepsy gene-testing panels. Understanding the pathophysiology of SRD is critical for developing targeted treatments. Recently-developed mouse models carrying human mutations of *SLC6A1* provide opportunities to understand SRD and test therapeutic interventions at the preclinical level.

Current preclinical studies ablate the *Slc6a1* gene to model SRD (Chiu et al., 2005; Gong et al., 2015; Shi et al., 2012; Chen et al., 2015). Mice lacking one copy of the *Slc6a1* gene have generalized epilepsy and no apparent behavioral deficits. Mice lacking both copies of *Slc6a1* have generalized epilepsy, motor deficits, anxiety-like behavior, and impaired performance on Morris water maze. Reports conflict on the pattern of locomotor activity, with findings of decreased or increased activity (Chiu et al., 2005; Gong et al., 2015; Chen et al., 2015). However, it is not clear whether all studies have used the same mutation or background strain, and it is uncertain to what extent some findings depend on sex. Whether ablation of *Slc6a1* models SRD is debated in light of recent studies that show that the majority (53/69) of disease-associated *SLC6A1* variants are missense mutations (Lal, 2021), leading to amino acid substitutions that may not be accurately recapitulated by a simple gene deletion. Here, we focus on a point mutation in the *SLC6A1* gene recently discovered in a child with epilepsy, developmental delay, sleep disturbances, muscle hypotonia, and attention deficits (Goodspeed et al., 2020; Mermer et al., 2021). This mutation results in retention of the GAT-1 protein in the endoplasmic reticulum, which may cause both reduced trafficking and toxicity to the endoplasmic reticulum (Mermer et al., 2021) We report results from a new mouse model in which the S295L missense mutation has been introduced at the homologous mouse *Slc6a1* locus. Using chronic wireless ECoG recordings, pharmacological manipulations, and behavioral assays, we evaluated whether this mouse might recapitulate key features of human SRD with the ultimate goal of understanding SRD pathophysiology and the implications for therapeutic development.

## RESULTS

### S295L variant confers sex-specific deficits in locomotor activity, muscle strength, and weight

Mice heterozygous for the patient-derived serine-to-leucine substitution at amino acid 295 of GAT-1 (GAT-1^S295L/+^) were engineered by introducing the corresponding DNA mutation into the mouse *Slc6a1* gene (see Methods). GAT-1^*S295L/+*^ mice were maintained with a heterozygote-by-wildtype mating scheme on a C57Bl/6J background, and offspring were fertile and viable, producing 50% male and female pups, ~7.36 pups per litter, in Mendelian ratios (**Figure 1A**). GAT-1^S295L/+^ females, but not males, weighed less than their GAT1^+/+^ sex-matched littermates (**Figure 1B**).

**Figure 1.**
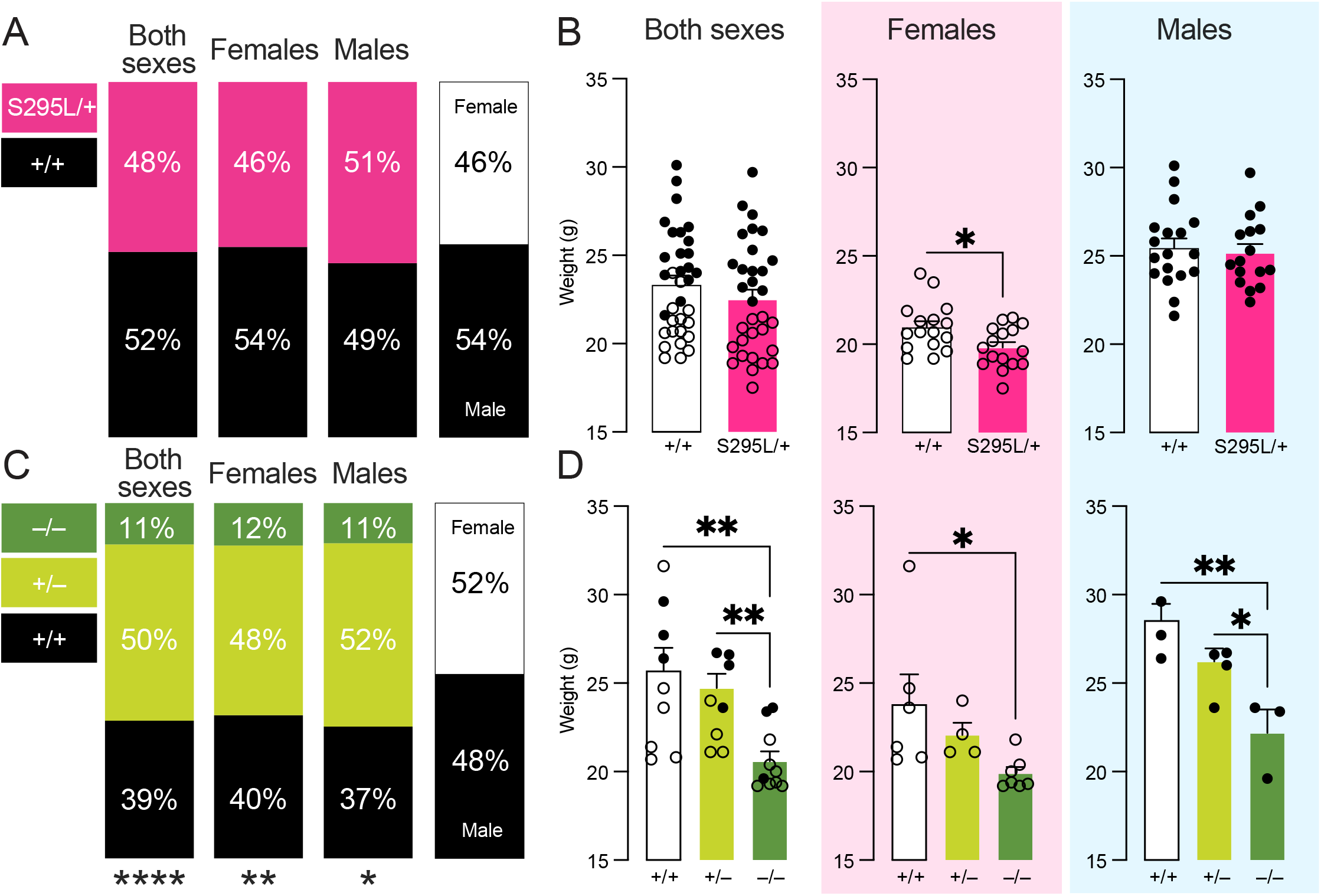
Mendelian ratio and body weight of GAT-1^S295L/+^ colony and GAT-1^+/−^ and GAT-1^−/−^ colony. **A.** Mendelian ratios are preserved in the GAT-1^S295L/+^ colony (expected 50% GAT-1^+/+^ and 50% GAT-1^S295L/+^, 1:1 Mendelian ratio). A total of 25 litters were obtained resulting in 184 mice (7.36 mice per litter) of both sexes: GAT-1^+/+^ n = 96, GAT-1^S295L/+^ n = 88 (Binomial test, two-tailed p=0.6059); 84 mice were female: GAT-1^+/+^ n = 45, GAT-1^S295L/+^ n = 39 (Binomial test, two-tailed p=0.5856); 100 mice were male: GAT-1^+/+^ n = 49, GAT-1^S295L/+^ n = 51 (Binomial test, two-tailed p=0.9204). Sex ratio was preserved (1:1 Mendelian ratio, Binomial test, two-tailed p=0.2688). **B.** Weight quantification of GAT-1^S295L/+^ mice. Pooled average of mice from implanted and non-implanted cohorts (left, t-test, p=0.2631; middle, t-test; right, t-test, p=0.6972). **C.** Mendelian ratios are skewed in the GAT-1^+/−^ colony (expected 25% GAT-1^+/+^, 50% GAT-1^+/−^, and 25% GAT-1^−/−^, 1:2:1 Mendelian ratio). A total of 23 litters were obtained resulting in 129 mice (5.6 mice per litter) of both sexes: GAT-1^+/+^ n = 50, GAT-1^+/−^ n = 64, GAT-1^−/−^ n = 15 (Chi-square 19.0, df = 2); 67 mice were female: GAT-1^+/+^ n = 27, GAT-1^+/−^ n = 32, GAT-1^−/−^ n = 8 (Chi-square 10.91, df = 2); 62 mice were male: GAT-1^+/+^ n = 23, GAT-1^+/−^ n = 32, GAT-1^−/−^ n = 7 (Chi-square 8.323, df = 2). Sex ratio was preserved (1:1 Mendelian ratio, Binomial test, two-tailed p=0.6612). **D.** Weight quantification in the knock-out cohort. (each one-way ANOVA, Tukey’s post-hoc test). * p<0.05, ** p<0.01. Each dot represents a single mouse (solid circles: males, open circles: females).

For comparison, we conducted key experiments in parallel in *Slc6a1* knock-out (–/–) mice (Jensen et al., 2003) and heterozygous (+/−) and wildtype (+/+) littermates, that we will refer to as GAT-1^−/−^, GAT-1^+/−^, and GAT-1^+/+^ mice, respectively. This colony of mice was maintained with a heterozygote-by-heterozygote mating scheme, also on a C57Bl/6J background. GAT-1^+/−^ mice were fertile and their pups were viable; they produced 50% male and female pups, ~5.6 mice per litter. However, ratios of offspring were non-Mendelian. GAT-1^−/−^ pups were under-represented, GAT-1^+/−^ normally represented, and GAT-1^+/+^ overrepresented in weanlings (**Figure 1C**). Female and male GAT-1^−/−^ mice weighed less than their GAT1^+/+^ sex-matched littermates (**Figure 1D**), consistent with prior reports (Chiu et al., 2005; Chen et al., 2015).

Behavior in GAT-1^−/−^ mice has previously been characterized by other researchers (Chiu et al., 2005; Gong et al., 2015; Shi et al., 2012; Chen et al., 2015), whose efforts we did not duplicate. To summarize briefly: GAT-1^+/−^ mice are reported to have no apparent behavioral deficits whereas GAT-1^−/−^ mice are reported to have motor deficits on rotarod, anxiety-like behavior, and impaired performance on Morris water maze. Reports conflict on the pattern of GAT-1^−/−^ locomotor activity, with findings of decreased or increased activity; it is not clear whether all groups used the same mutation and background strain. Therefore, we evaluated locomotor activity in the home cage in the GAT-1 knock-out colony to resolve this discrepancy. We undertook more detailed behavioral characterization in the GAT-1^S295L/+^ mice, which, to our knowledge, have not been previously characterized.

GAT-1^S295L/+^ mice demonstrated no anxiety-related behavior, as assessed with the elevated plus maze (**Supplemental Figure 1**), light-dark box (**Supplemental Figure 2**), and time spent in the center of the open field (OF) test (**Supplemental Figure 3,4**). Working memory was also preserved, as measured by the Y-maze (**Supplemental Figure 5**). Social behavior, assessed with the three-chamber sociability assay and social dominance tube test, was found to be normal in GAT-1^S295L/+^ mice (**Supplemental Figure 6**). While the main outcomes of each of these tasks did not exclude the null hypothesis, embedded measures of locomotor activity within each behavioral task showed small but persistent differences by genotype in female mice but not in male mice (**Figure 2A-E**). Of note, no effect was seen during initial OF testing (**Supplemental Figure 3**), the first assessment in the series. To account for the effects of habituation on vigilance state, we repeated OF at the end of the testing battery, and, under these conditions, female GAT-1^S295L/+^ mice traveled less distance (**Figure 2A-B**).

**Figure 2.**
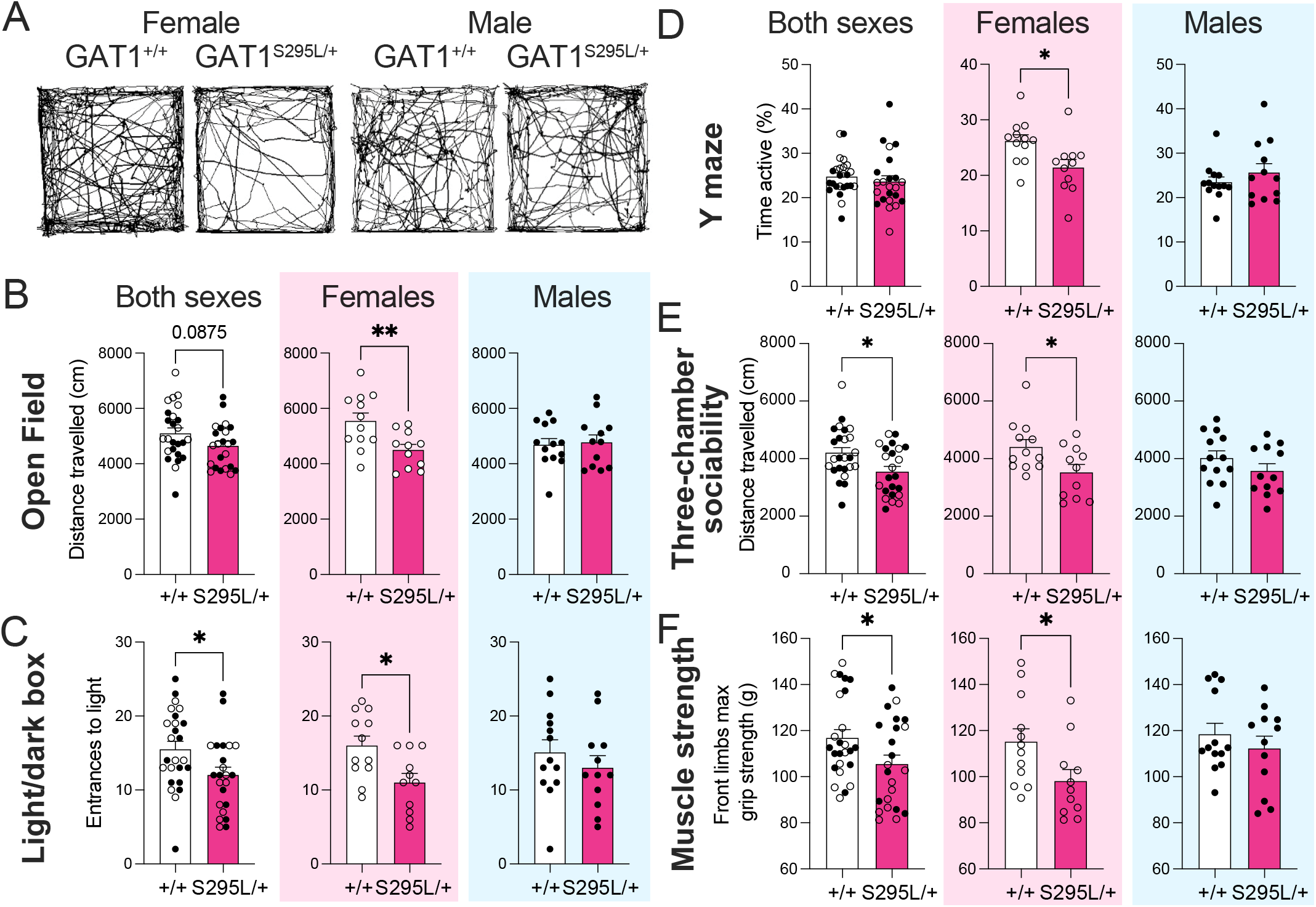
GAT-1^S295/+^ female but not male mice exhibit decreased locomotion and muscle grip strength. **A.** Representative locomotion tracks during 10 minutes of exploratory activity on the second day in the open field (see Supplemental Figures 2,3 for a detailed analysis of the open field performance). **B.** Total distance travelled (left, unpaired t-test, p=0.0875, n=25,23; middle, unpaired t-test, p=0.0078, n=12,11; right, unpaired t-test, p=0.7926, n=13,12; see Supplemental Figures 2,3 for a detailed analysis of the open field analyses). **C.** Entrances to light chamber (left, unpaired t-test p=0.0235, n=25,23; middle, unpaired t-test, p=0.0105, n=12,11; right, unpaired t-test, p=0.3890, n=13,12). **D.** Time active in the Y maze (left, unpaired t-test, p=0.4578, n=25,23; middle, unpaired t-test, p=0.0157, n=12,11; right, unpaired t-test, p=0.3427, n=13,12). **E.** Distance travelled in the three-chamber sociability test (left, unpaired t-test, p=0.0118, n=25,23; middle, unpaired t-test, p=0.0258, n=12,11; right, unpaired t-test, p=0.21, n=13,12). **F.** Forelimb maximum grip strength (left, unpaired t-test, p=0.0361, n=25,23; middle, unpaired t-test, p=0.0348, n=12,11; right, unpaired t-test, p=0.3938, n=13,12; see Supplemental Figure 1 for a detailed analysis of muscle grip strength). * p<0.05, ** p<0.01. Each dot represents a single mouse (solid circles: males, open circles: females).

Results from behavioral assessments suggested a motor deficit in female GAT-1^S295L/+^ mice. Therefore, we further characterized motor performance with a measure of muscle strength. Female GAT1^S295L/+^ mice showed a significant reduction in forelimb grip strength compared to wildtype littermates (**Figure 2F**).

In summary, our noninvasive observations revealed a phenotype in female GAT-1^S295L/+^ mice, namely decreased locomotor activity, muscle strength, and body weight.

### The S295L variant confers an epileptic phenotype including nonconvulsive seizures characterized by SWDs and motor arrest

As seizures are a common feature of SRD (Johannesen et al., 2018), we hypothesized that the GAT-1^S295L/+^ variant would confer a tendency towards epilepsy in mice. To evaluate this possibility, we obtained chronic wireless ECoG and electromyographic (EMG) recordings from GAT-1^S295L/+^ mice and wild-type GAT-1^+/+^ littermate controls. During 48 hours of baseline recording, we observed frequent SWDs in GAT-1^S295L/+^ mice but not in GAT-1^+/+^ littermate controls (**Figure 3**). SWDs were associated with motor arrest (**Figure 3A-B**) and mild head drop consistent with myoclonic atonic epilepsy (Oguni et al., 1992). Baseline ECoG power, assessed by root mean square (RMS) amplitude sampled during a spike-free interval, was similar in GAT-1^S295L/+^ and GAT-1^+/+^ mice (**Figure 3C**). An increased burden of spikes and SWDs was apparent during both diurnal and nocturnal periods (**Figure 3D-G**). Male and female GAT-1^S295L/+^ mice had comparable burdens of spikes (light period p=0.66; dark period p=0.34; n=3,4; unpaired t-tests) and SWDs (light period p=0.10; dark period p=0.08; n=3,4; unpaired t-tests).

**Figure 3.**
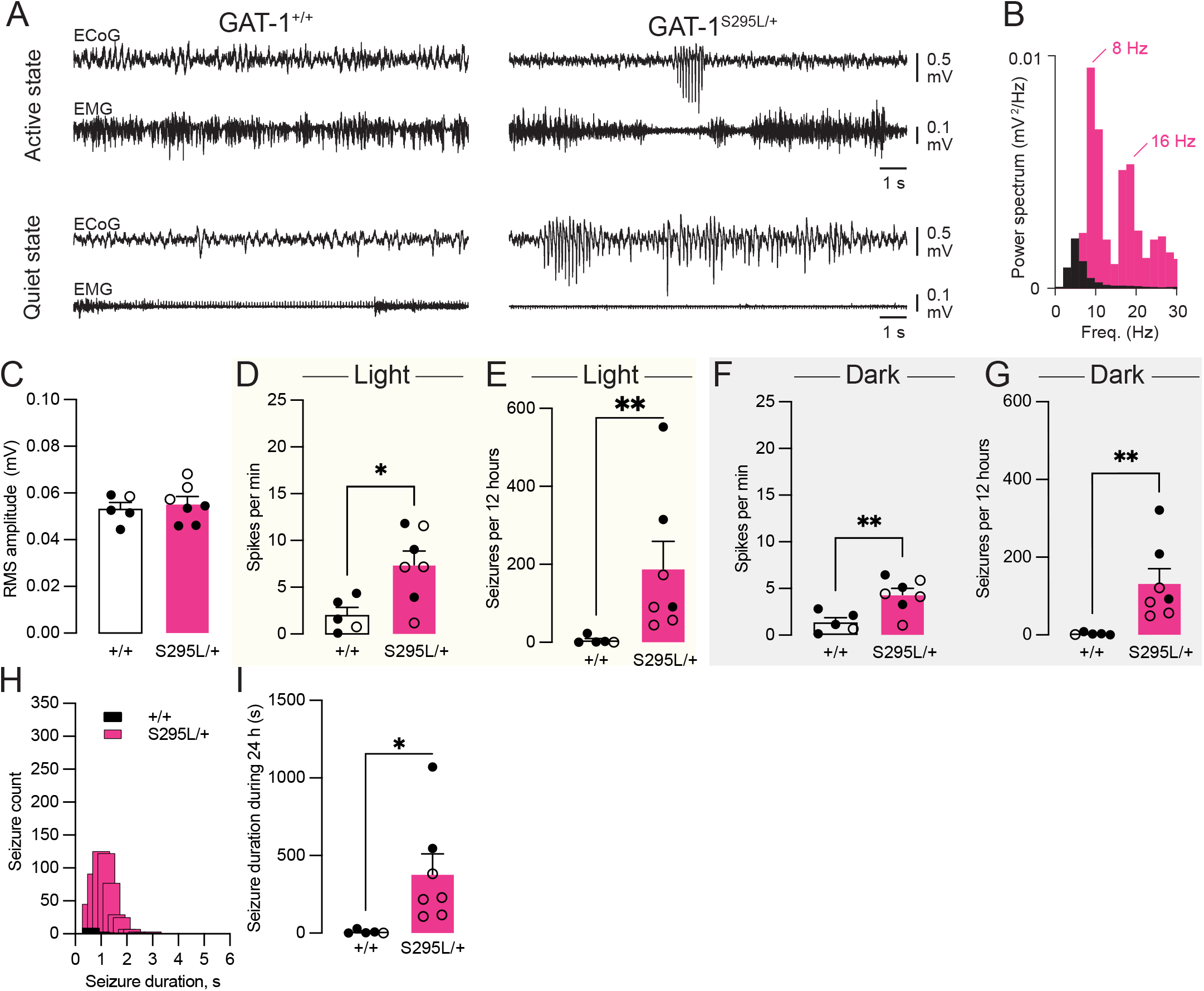
GAT-1^S295/+^ mice have SWDs associated with motor arrest. **A.** ECoG and EMG recordings from a representative wild-type and S295L variant mouse during active and quiet behavioral states. During active behavior (top traces), note the interruption of muscle activity observed in the EMG recording during an SWD in a GAT-1^S295L/+^ mouse. Also note the frequent SWDs associated with lack of EMG activity during quiet behavioral states (bottom traces). **B.** Spectral density plot of ECoG, evaluated during the active segment shown in (A). Note a peak power at 8Hz and harmonic at 16Hz in GAT-1^S295L/+^ (pink), compared with weaker power peak at ~4Hz in wild-type GAT-1^+/+^ (black). **C.** Baseline power, quantified as RMS amplitude sampled during a spike-free interval during active wakefulness (p=0.62 unpaired t-test, n=5,7). **D.** Spike counts per minute during the day, detected as events greater than 7-fold the RMS amplitude of baseline awake ECoG, and averaged over two consecutive 12-h light periods (n=5,7, unpaired t-test). **E.** Quantification of SWDs during the light period two consecutive 12-h dark periods (n=5,7, unpaired t-test). **G.** Quantification of SWDs during the dark period (Mann-Whitney test, n=5,7). **H.** Histogram of the incidence and duration of SWDs. **I.** Cumulative time spent seizing over a 24h period (Welch’s t-test, n=5,7). * p<0.05, ** p<0.01. Each dot represents a single mouse (solid circles: males, open circles: females).

Both the number and duration of SWDs (**Figure 3H**) were substantial in GAT-1^S295L/+^ mice (grand mean 1.19 ± 0.01s from 2253 events in 7 GAT-1^S295L/+^ mice), with a cumulative seizure burden of 6.4 ± 2.1 min in a representative 24h interval (**Figure 3I**).

To evaluate contributions of gene dosage to observed ECoG effects without toxicity from endoplasmic reticulum trafficking defects (Mermer et al., 2021), we performed recordings in mice lacking either one or two copies of GAT-1 (Jensen et al., 2003; Chiu et al., 2005).

ECoG recordings in GAT-1-deficient mice demonstrated a dose-response relationship between the number of *Slc6a1* alleles ablated and the severity of the ECoG phenotype. Specifically, GAT-1^+/−^ mice had frequent SWDs, while GAT-1^−/−^ had a very severe ECoG phenotype with episodes of continuous SWDs corresponding to prolonged periods of immobility (**Figure 4A,B**). ECoG baseline RMS amplitude, sampled from a spike-free interval, was significantly increased in GAT-1^−/−^ mice compared with GAT-1^+/−^ and wildtype GAT-1^+/+^ littermates (**Figure 4C**). While SWD duration in GAT-1^−/−^ mice was underestimated by objective measures (long-episode SWDs were truncated given that some spike-wave complexes went undetected by a stringent definition of spike threshold), quantification of spike burden (**Figure 4D–F**) as well as number and duration of detected SWDs (**Figure 4E, G,H**) showed a gradient of ECoG abnormality with increasing GAT-1 deficiency. Compared with males, females had similar ECoG abnormalities within affected genotypes (GAT-1^+/−^ spikes per min light period p=0.78, dark period p=0.80; Seizures per 12 hours p=0.054, n=5,4; GAT^−/−^ spikes per min light period p=0.83, dark period p=0.83; Seizures per 12 hours p=0.74, n=4,5, unpaired t-tests, **Figure 4D-G**). The cumulative duration of SWDs in GAT^+/−^ mice reached 3.6 ± 1.4 min whereas in GAT^−/−^ mice this was increased to 24.6 ± 5.1 min in a 24h interval (**Figure 4I**).

**Figure 4.**
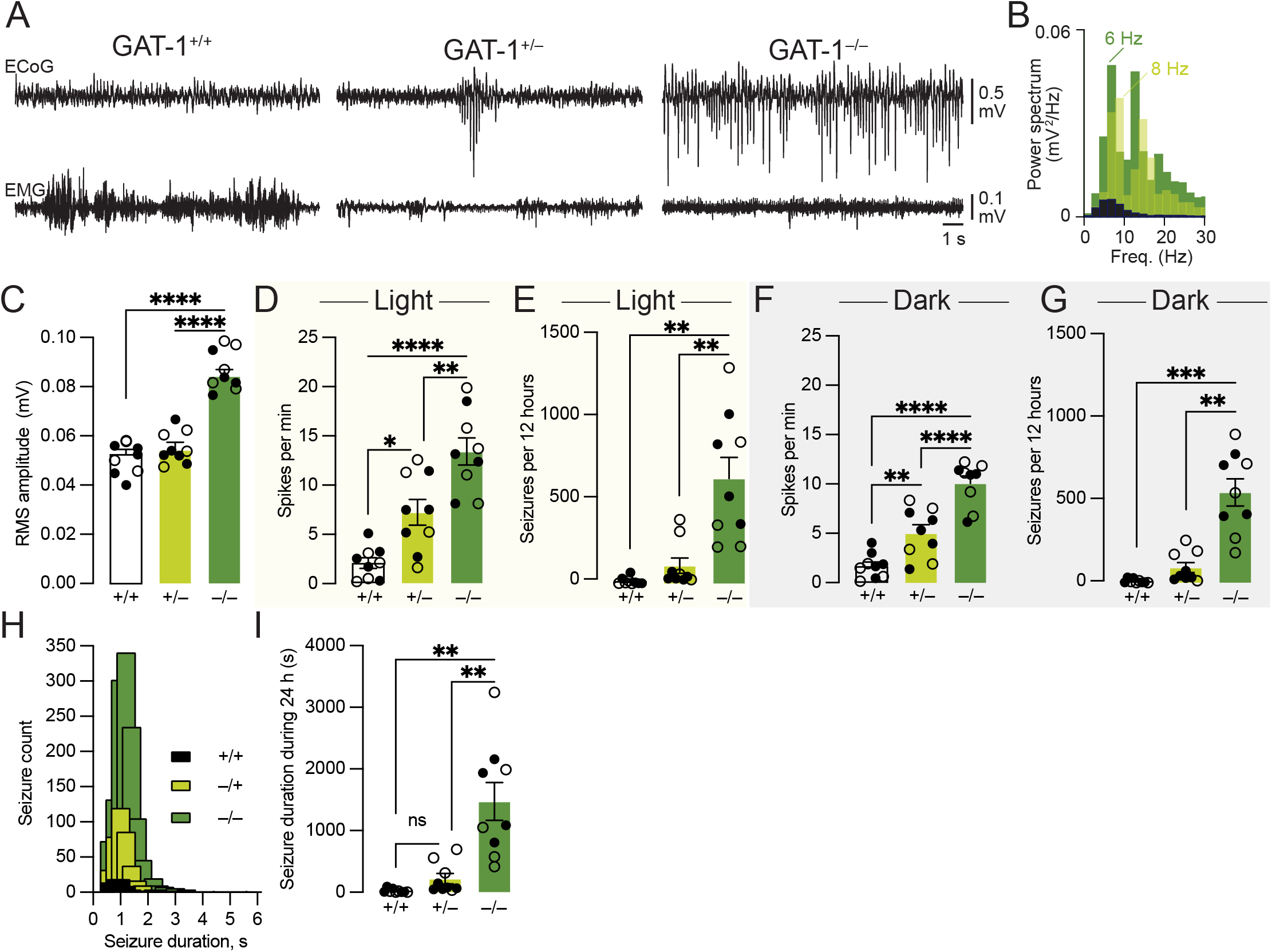
GAT-1 gene dosage correlates with the severity of ECoG abnormalities. **A.** Simultaneously recorded ECoG and EMG activity from from representative mice. During awake unrestricted activity in the GAT-1^+/+^ mouse (left traces), low amplitude desynchronized ECoG activity (first row) is accompanied by high amplitude spontaneous EMG activity (second row). In the GAT-1^+/−^ mouse (middle traces), the SWD is associated with a pause of voluntary motor activity seen as decrement in EMG amplitude. In the GAT-1^−/−^ mouse, ECoG is dominated by continuous SWDs associated with immobility indicated by quiescent EMG. **B**. Spectral density plot of ECoG during the interval shown in (A). Note a peak power at 6Hz and a harmonic at 12Hz in GAT-1^−/−^ (dark green), compared with a peak at ~8Hz and harmonic at 16Hz in GAT-1^+/−^ (light green). **C.** Baseline power, quantified as RMS amplitude sampled during a spike-free interval during active wakefulness (ANOVA p<0.0001; Tukey’s multiple comparisons; n=9,9,9). **D.** Spike quantification, averaged over two consecutive 12-h light periods (ANOVA p<0.0001, Tukey’s multiple comparisons, n=9,9,9). **E.** SWD quantification, assessed during the light period (Brown-Forsythe p<0.0005, Dunnett’s multiple comparisons, n=9,9,9). **F.** Quantification of spikes averaged over two consecutive dark periods (ANOVA p=0.04, Tukey’s multiple comparisons, n=9,9,9.) **G.** SWD quantification, assessed during the dark period (Brown-Forsythe p<0.0001, Dunnett’s multiple comparisons, n=9,9,9). **D-G**. Compared with males, females had similar ECoG abnormalities within affected genotypes (GAT-1^+/−^ spikes per min light period p=0.78, dark period p=0.80; Seizures per 12 hours p=0.054, n=5,4; GAT^−/−^ spikes per min light period p=0.83, dark period p=0.83; Seizures per 12 hours p=0.74, n=4,5, unpaired t-tests). **H.** Histogram plotting incidence and duration of SWDs (GAT-1^+/+^ black, GAT-1^+/−^ light green, GAT-1^−/−^ dark green). **I.** Cumulative seizure duration summated over 24h (Brown-Forsythe p=0.0006; Dunnett’s multiple comparisons, n=9,9,9). *p<0.05, **p<0.01, ***p<0.001, ****p<0.0001. Each dot represents a single mouse (solid circles: males, open circles: females).

While we did not directly compare GAT-1^S295L/+^ to GAT-1^+/−^ mice, qualitatively, GAT-1^S295L/+^ had an intermediate phenotype along this continuum, suggesting some residual GAT-1 function from the intact allele.

### Electrocorticographic abnormalities correlate with functional deficits in nesting and locomotion

Next, we asked whether ECoG abnormalities might intrude upon the awake state to such a degree that they interfere with performance of activities of daily living. Mice are nocturnal mammals, active during the dark period and sleeping predominantly during the light period (Refinetti, 2004). Locomotor activity was assessed continuously throughout both light and dark periods by wireless telemetry system during baseline ECoG recordings. During the dark period, activity was decreased in GAT^−/−^ mice compared with littermate controls (**Figure 5A**), in agreement with a previous report that found that home cage activity was reduced in these mice (Chiu et al., 2005). By contrast, GAT-1^S295L/+^ (**Figure 5A**), and GAT-1^+/−^ mice showed levels of activity comparable to GAT-1^+/+^ littermates (**Figure 5B**). As expected, activity was higher in all genotypes during the dark period compared with the light period (data not shown). Activity in the light period did not differ significantly in GAT-1^S295L/+^, GAT-1^+/−^, nor GAT-1^−/−^ mice compared with GAT-1^+/+^ littermate controls (data not shown).

**Figure 5.**
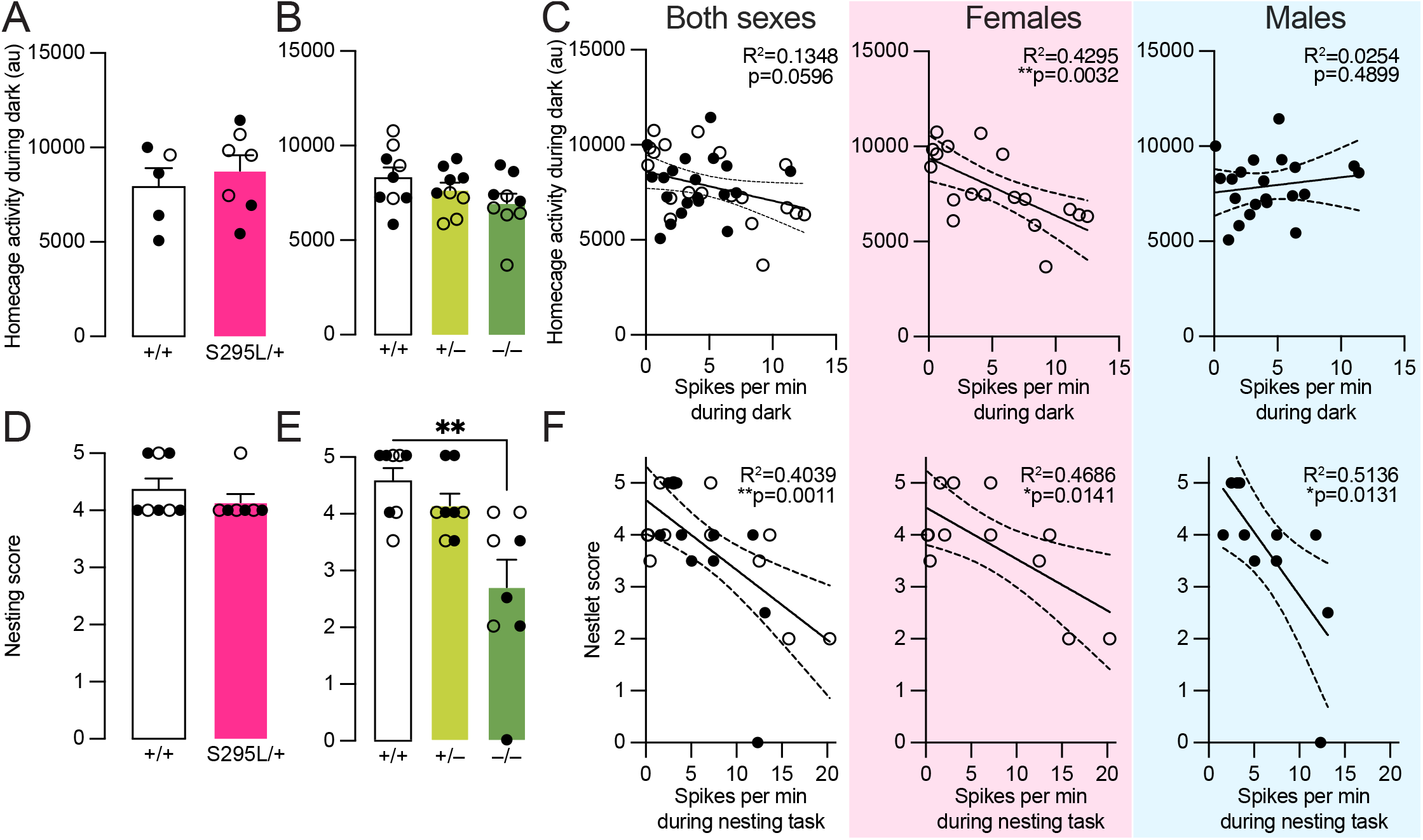
Electrocorticographic features predict behavioral outcomes across genotypes. **A.** Nocturnal activity counts, measured as the average of two consecutive 12h dark periods in the home cage (unpaired t-test, p=0.5323, n=5,7). **B.** Nocturnal activity counts (ANOVA p=0.1460, n=9,9,9). **C.** Spikes measured during the nocturnal period predicted locomotor activity over the same duration, in aggregate (left, n=39). Note that the relationship was driven by female subjects (n=19), and no relationship was seen in male mice (n=20). For analyses shown in C,F, all 5 genotypes from both colonies were included (GAT-1^S295L/+^ (n=5) and GAT-1^+/+^ littermates (n=7), GAT-1^+/−^ (n=9), GAT-1^−/−^ (n=9), and GAT-1^+/+^ littermates (n=9)). Correlation coefficient calculated by Pearson’s R; linear regression (solid lines) with 95% confidence intervals (dashed lines). **D.** Nest building score in the S295L colony; scored at 48 hours on a 6-point (0-5) scale. This panel captures data from some mice without ECoG or locomotor activity (underwent device implantation surgery and recovery but no ECoG was recorded due to transmitter failure). Unpaired t-test, p=0.3457, n=8,7. **E.** Quantification of nesting behavior in the knock-out colony (ANOVA; post-hoc t-tests with Dunnett’s correction, n=8,8,8). **F.** Spikes measured during the 48-hour nesting assessment (expressed as an average rate) predicted nestlet quality, in aggregate (left), and in both female (middle) and male mice (right). Pearson’s correlation coefficient; linear regression (solid lines) with 95% confidence intervals (dashed lines). * p<0.05, ** p<0.01. Each dot represents a single mouse (solid circles: males, open circles: females).

We next evaluated whether there was a generalizable relationship between an individual subject’s locomotor activity and ECoG activity, regardless of genotype. Indeed, in pooled analysis incorporating all 5 genotypes from both the S295L colony and the GAT-1 knockout colony, spike burden was negatively correlated with locomotor activity measured during the same time interval. The inverse relationship between spike count and locomotor activity was seen during the dark (active) period (**Figure 5C**) but not during the light (sleep) period, and only applied to female mice (**Figure 5C**). The female bias of this effect fits well with our observation that females but not males in the non-implanted behavioral cohort exhibited abnormalities in locomotor activity and further suggests that those subjects with more pronounced locomotor hypoactivity were those with more severe ECoG abnormalities.

Next, we evaluated nest building behavior concurrent with baseline physiology recordings. GAT-1^S295L/+^ mice constructed nests indistinguishable from GAT-1^+/+^ littermates (**Figure 5D**). By contrast, GAT-1^−/−^ mice constructed poor quality nests compared with GAT^+/+^ littermates (**Figure 5E**). As hypothesized, nesting performance correlated with the severity of ECoG abnormalities as quantified by spike burden during the nest-building opportunity (**Figure 5F**). Both male and female mice demonstrated a negative correlation between spike burden and nest quality (**Figure 5F**). In the S295L colony, we found no effect of genotype on nesting, in aggregate or by sex (24h GAT-1^+/+^ vs GAT-1^S295L/+^ p=0.347, n=24,23, unpaired t-test; GAT-1^S295L/+^ male vs female p=0.75, n=12,11, unpaired t-test; 48h GAT-1^+/+^ vs GAT-1^S295L/+^ p=0.38, p=24,23, unpaired t-test; GAT-1^S295L/+^ male vs female p>0.999, n=12,11).

Results shown here suggest that the ECoG spike burden accounted for ~30% of the variability in nesting (in females) and ~40–50% of the variability in locomotor activity (in both males and females). This result links the severity of epileptic abnormalities with ethologically relevant activities of daily living.

### Seizures are bi-directionally modulated by NO-711 and ethosuximide

Following completion of 48 hour baseline physiology recordings, we evaluated the ECoG-device-implanted cohort of mice pharmacologically using a within-subject repeated measures design. To assess residual GAT-1 function, we tested ECoG responses to acute application of the GAT-1 antagonist NO-711. Furthermore, we evaluated ECoG responses to the anti-epileptic drug ethosuximide.

Compared to vehicle, systemic NO-711 acutely provoked SWDs in GAT-1^+/+^ mice, and dramatically increased the incidence of SWDs in GAT-1^S295L/+^ mice, inducing nearly continuous SWDs (**Figure 6**) reminiscent of long episodes seen in GAT-1^−/−^ mice (**Figure 4**). Ethosuximide 200 mg/kg, by contrast, eliminated spontaneous SWDs in GAT-1^S295L/+^ mice (**Figure 6A**). Drug effects were robust in both light and dark periods (**Figure 6B-E**).

**Figure 6.**
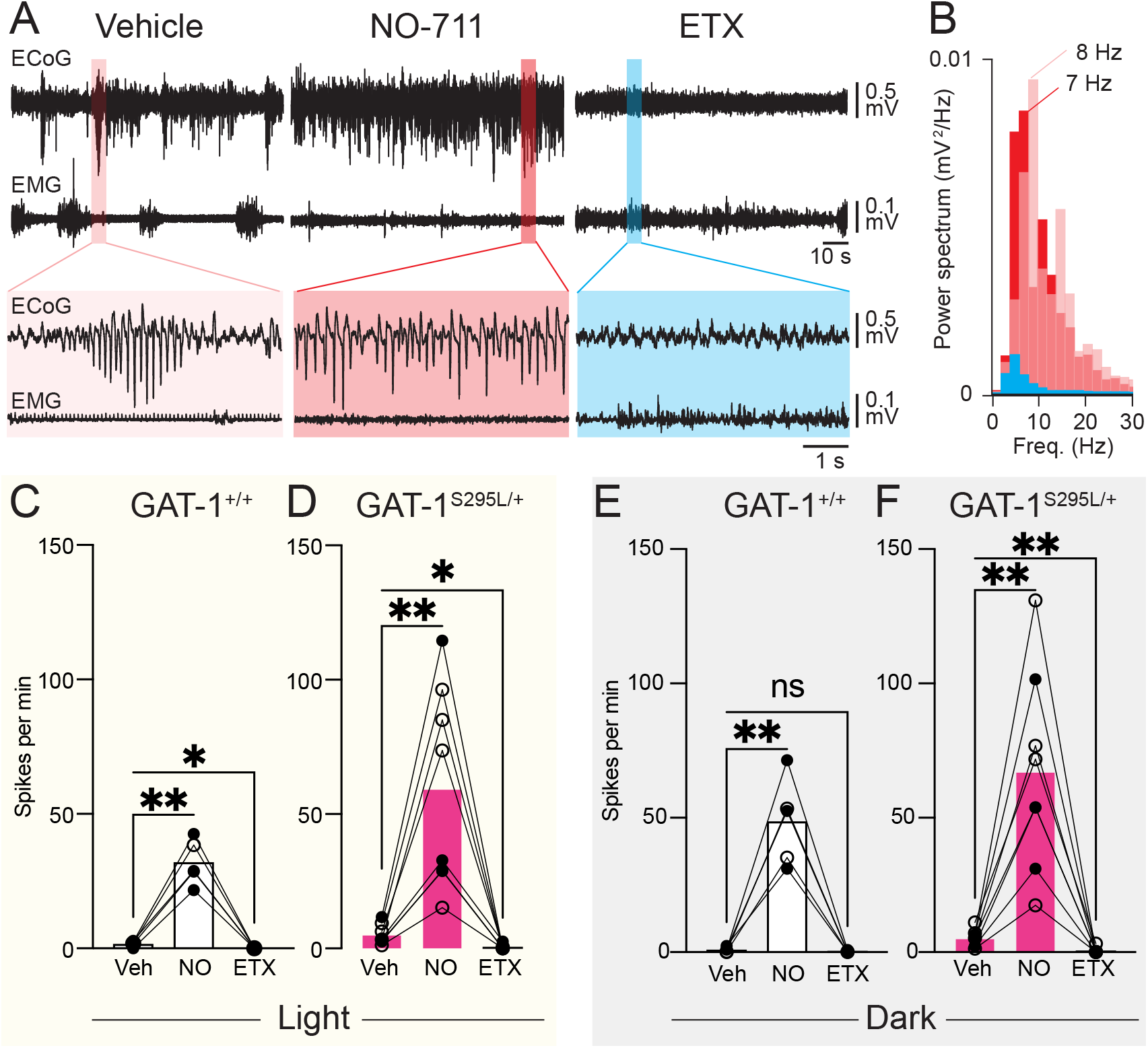
Pharmacosensitivity of GAT-1^S295L/+^ mice. **A.** ECoG and EMG traces from a representative GAT-1^S295L/+^ mouse. Note the spontaneous SWDs under control conditions (injection of normal saline 0.01 ml/g body weight) increased by GAT-1 antagonist NO-711 (10 mg/kg i.p.) and eliminated by ethosuximide (ETX, 200 mg/kg i.p.). There was a 48h wash-out between each treatment. **B.** Power frequency histogram from the expanded time base segments shown in (A). Under vehicle conditions, note peak at 8Hz (pink) and harmonic at 16Hz. During treatment with NO-711, the peak shifts to 7Hz with a harmonic at 14Hz (red). During treatment with ethosuximide, power decreases and peak shifts to ~4Hz (blue). **C-F**. Population data summarizing ECoG changes in response to vehicle, NO-711, and ethosuximide. Data are sampled from the first hour post-injection, at the start of the light (B,C) and dark periods (D,E). The sequence of drug administration (NO-711 vs ethosuximide) was alternated in interleaved animals to control for order effects, if any. RM-ANOVA with Dunnett’s multiple comparisons tests, n=5,8. * p<0.05, ** p<0.01. Each dot represents a single mouse (solid circles: males, open circles: females).

Concordant results were seen in GAT-1 deficient mice (**Figure 7**). Thus, in GAT-1^+/−^ mice, NO-711 increased and ethosuximide eliminated SWDs during diurnal and nocturnal periods (**Figure 7 B,E**). NO-711 was effective at provoking SWDs in GAT-1^+/+^ littermates during both day and night (**Figure 7 A,C**). By contrast, NO-711 had no consistent effect on GAT-1^−/−^ mice, an important control as these mice lack the target of the drug (**Figure 7 C,F**). Consistent with the severe ECoG phenotype in GAT-1^−/−^ mice, SWDs were resistant to ethosuximide at the dose and time-point tested in knock-out mice.

**Figure 7.**
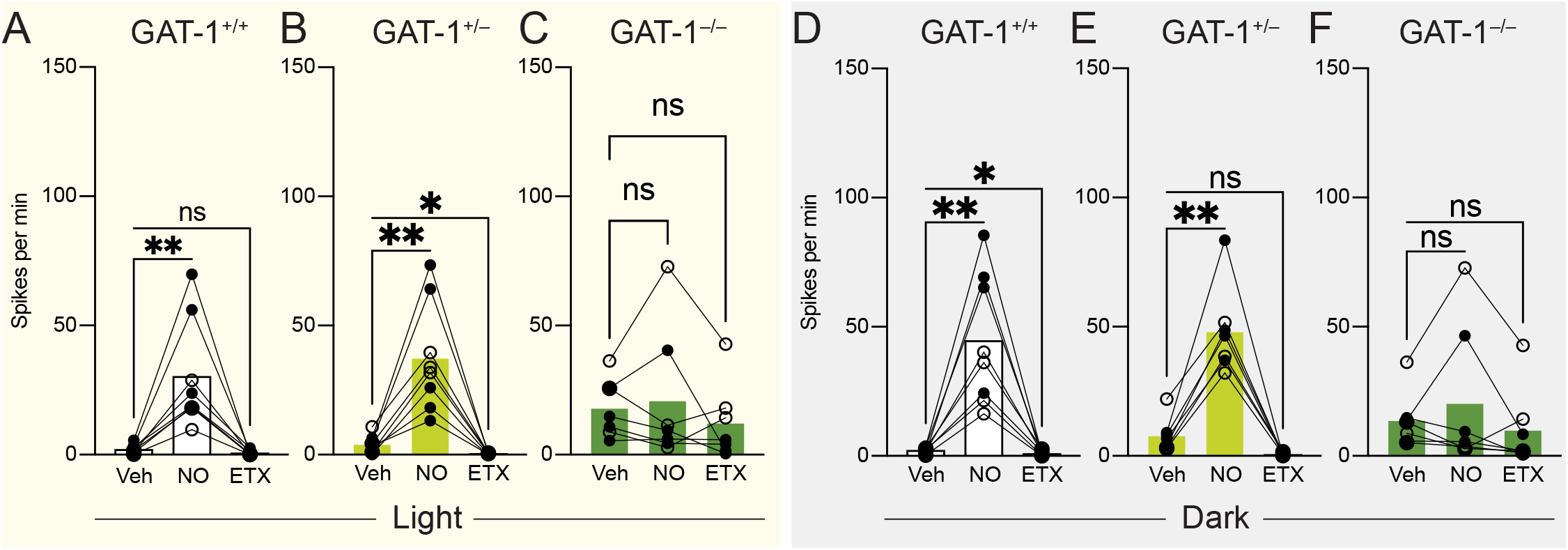
Pharmacosensitivity of GAT-1 knock-out mice. **A-C.** Population data plotting spike burden under conditions of vehicle, NO-711, and ethosuximide administration during the light period (A. RM-ANOVA p=0.005; Dunnett’s multiple comparison test; n=8. B. RM-ANOVA p=0.002; Dunnett’s multiple comparisons; n=8); C. RM-ANOVA p=0.37, n=8). **D-F**. Population data plotting spike burden under conditions of vehicle, NO-711, and ethosuximide administration during the dark period (D. RM-ANOVA p=0.002, Dunnett’s multiple comparisons, n=8; E. RM-ANOVA p=0.0001, Dunnett’s multiple comparisons, n=7; F. RM-ANOVA p=0.026, n=7). Data shown represent the first hour post-injection, at the start of the light (A,B,C) and dark periods (D,E,F). The sequence of drug administration (NO-711 vs ethosuximide) was alternated in interleaved animals to mitigate any order effects, similar to Fig. 6. * p<0.05, ** p<0.01. Each dot represents a single mouse (solid circles: males, open circles: females).

In summary, GAT-1^S295L/+^ mice had residual intact GAT-1 function demonstrated by a marked ECoG response to NO-711, at a dose that did not affect GAT-1^−/−^ mice. SWDs were sensitive to ethosuximide in GAT-1^S295L/+^ and GAT-1^+/−^ mice but not in the most severely affected mice (GAT-1^−/−^). These pharmacological observations support the hypothesis that S295L leads to a partial loss of GAT-1 function.

## DISCUSSION

Introducing the GAT-1^S295L/+^ mutation in mice recapitulated major features of human SRD, including absence or atonic seizures and behavioral deficits that correlate with the severity of ECoG abnormalities during the awake, active period. Our data support the general model that this mutation leads to loss of function of the gene product, and that its functional impact results from insufficient surface expression of GAT-1.

### S295L faithfully recapitulates human SRD features

Our finding that introducing the S295L variant in the *Slc6a1* gene resulted in SWDs with motor/behavioral arrest and mild head drop is consistent with absence and myoclonic atonic epileptic seizures (Oguni et al., 1992), and reduced nesting behavior. Although both sexes showed a similar seizure phenotype, only female mice showed reduced muscle strength and locomotion; and the spike burden was correlated with reduced activity in female mice. As homozygous knock-out (GAT^−/−^) mice had previously been characterized in several studies (Chiu et al., 2005; Gong et al., 2015; Shi et al., 2012; Chen et al., 2015), we did not undertake a detailed behavioral assessment except where it informed our interpretation of ceiling and floor effects of GAT-1 function. Of note, we did find decreased locomotor activity in GAT-1^−/−^ mice in their the home cages in agreement with Chiu and colleagues, thereby helping to resolve conflicting reports in the literature of increased versus decreased locomotor activity in these mice (Chiu et al., 2005). Additionally, we observed a nesting deficit in GAT^−/−^ mice which has not (to our knowledge) been previously reported.

The finding that seizure burden was correlated with ethological behavioral outcomes raises the possibility that electrocorticographic abnormalities may intrude upon the awake state to such a degree that they interfere with the performance of activities of daily living such as ambulation and nesting and feeding. As with other developmental epileptic encephalopathies, a key question is whether reducing seizure burden would rescue the behavioral phenotype. On the one hand, mutations in the *Slc6a1* gene may cause developmental circuit abnormalities independent of seizures. In that case, correcting seizures would not correct the underlying disorder of development. Indeed, during brain development, GAT-1 is known to mediate GABA release and signaling, which is neurotrophic and is involved in a variety of processes from cell proliferation and migration to synapse formation and plasticity (Ben-Ari, 2002). On the other hand, if seizures intrude on the performance of a behavioral task in the moment, successfully treating the seizures should rescue performance. Another possibility is that seizures occur during a critical period of development, representing an opportunity cost; a brain of a child with frequent seizures may not able to learn age-appropriate tasks and build experience-dependent circuits. The S295L mouse provides an excellent model to begin addressing such questions.

### Is S295L a simple loss of function? Implications for treatments

Most pathogenic mutations in the *SLC6A1* gene result in partial or complete loss of GAT-1 function (Mermer et al., 2021), which is consistent with the finding that GAT-1^S295L/+^ mice showed SWDs qualitatively similar to those seen in GAT-1^+/−^ heterozygous mice lacking one copy of the gene. However, GAT-1^+/−^ mice have a mild form of typical absence epilepsy and no behavioral deficits (Chiu et al., 2005), whereas GAT-1^S295L/+^ mice show abnormalities in muscle strength and locomotion, suggesting a toxic effect of the point mutation. These findings are in agreement with the fact that S295L mutation results in retention of the protein in the endoplasmic reticulum in both human and mouse cells, which could cause toxicity and/or partial retention of the normal copy of the protein as well (Mermer et al., 2021). Our finding that systemic treatment with GAT-1 blocker NO-711 enhanced seizures in GAT-1^S295L/+^ mice suggests that there is residual GAT-1 protein expression in these mice. We conclude that although knockout mice have been a useful in showing that loss of function of GAT-1 could lead to generalized absence epilepsy, SRD is a more complex syndrome that warrants further investigation.

If the S295L variant confers pathogenicity by a mechanism of haploinsufficiency, gene therapy strategies that boost expression from the native allele might be successful. If, on the other hand, the S295L variant acts as a dominant negative, (i.e. by interfering with translation or trafficking of GAT-1 from the intact allele), therapeutic strategies to chaperone the folding and transport of the normal protein, or ablating the mutant allele, may be necessary. Future studies will test these hypotheses.

### Role of GAT-1 in epilepsy: potential mechanisms

Prior to the identification of *SLC6A1* mutations in epilepsy patients in 2015 (Carvill et al., 2015), studies had demonstrated GAT-1 dysfunction in rodent models of generalized absence epilepsy (Cope et al., 2009). Furthermore, the expression of GAT-1 is reduced around dentate granule cells in resected hippocampal tissue from patients with temporal lobe epilepsy (Lee et al., 2006). While the mechanisms leading to SRD are unknown, several lines of evidence from studies from knockout mice point to tonic GABA current as a potential mediator of absence seizures (Cope et al., 2009). The model that emerged from the studies listed above is that GAT-1 dysfunction reduces GABA uptake, leading to increased extrasynaptic tonic GABA current in the postsynaptic neuron. Consistent with this model, a tonic current mediated by the extrasynaptic GABA_A_ receptor is increased in thalamocortical neurons in absence epilepsy (Cope et al., 2009). Our studies confirm that a single missense mutation – shown to retain GAT-1 in the endoplasmic reticulum resulting in less GAT-1 at the membrane (Mermer et al., 2021) – is sufficient to cause SWDs, in agreement with findings from mice with deletion of one copy of *Slc6a1*. Unexpectedly, the point mutation resulted in reduced locomotion and muscle strength, which have not been observed with a simple deletion of one copy of *Slc6a1*. We propose that S295L mice recapitulate SRD better than mice with ablation of *Slc6a1* and therefore is a better model for SRD.

### Importance of sex and genetic background in behavioral assays

The sex-specific behavioral phenotype we discovered here was an unexpected result. Although it remains to be determined why female mice are more functionally impacted than male mice despite similar overall seizure burden of seizures, our findings highlight the importance of evaluating phenotypes and reporting results carefully by sex in the preclinical epilepsy field. We acknowledge that apart from sex, other factors likely influence the penetrance and expression of disease. Genetic context may matter; the phenotype seen on C56Bl/6J background may differ in other inbred strains or in outbred populations. Another possibility is that there are sex-specific compensations by other GABA transporters that could explain the lack of certain behavioral deficits in GAT-1^S295L/+^ males. For instance, in the case of temporal lobe epilepsy, GAT-1 deficits can be accompanied by GAT-3 upregulation in sclerotic hippocampi, suggesting compensatory mechanisms that may be a key feature of resilience in certain cells/regions (Lee et al., 2006). Future studies will investigate sex-specific mechanisms of SWDs and behavioral deficits, and whether these two are caused by the same circuits.

### Broader implications beyond genetic neurodevelopmental disorders

GAT-1 is an emerging target in neurological disorders other than SRD. Reduced levels of GAT-1 have been reported in patients with temporal lobe epilepsy and stroke, in which the *SLC6A1* gene is not a reported susceptibility gene. Furthermore, GAT-1 may serve as a novel therapeutic target for stroke treatment during the repair phase (Clarkson et al., 2010). Therefore, in addition to benefiting the individuals impacted by *SLC6A1* mutations, understanding the role of GAT-1 in brain function has the potential to benefit individuals suffering from neurological disorders such as stroke, which is a major cause of disability, and a common cause of epilepsy (Paz et al., 2013; Sarecka-Hujar and Kopyta, 2019; Myint et al., 2006).

## MATERIALS AND METHODS

### Data availability

The data that support the findings of this study are available from the corresponding author upon reasonable request.

### Animals

We performed all experiments per protocols approved by the Institutional Animal Care and Use Committee at the University of California, San Francisco and Gladstone Institutes, with institutional oversight. Experiments were conducted according to ARRIVE guidelines (Kilkenny et al., 2012) and recommendations to facilitate transparent reporting (Landis et al., 2012). Researchers were blinded to experimental groups during data collection and analysis. All biological variables were documented. Littermate controls were used for each experiment: GAT-1^S295L/+^ mice were compared with their GAT-1^+/+^ littermate controls, and knock-out GAT-1^+/−^ and GAT-1^+/−^ mice were compared with their GAT-1^+/+^ littermate controls. Balanced numbers of male and female mice were assigned to each experimental group. Precautions were taken to minimize distress and the number of animals used in each set of experiments. Mice were bred on an isogenic C57BL/6J background, housed in a pathogen-free barrier facility on a standard 12-hour light/ dark cycle with ad libitum access to food and water. Mice were genotyped from tail tissue collected at weaning and after death using Transnetyx genotyping services.

### GAT-1^S295L/+^ mice

GAT-1^S295L/+^ mice were obtained from Shanghai Model Organisms (C57BL/6-*Slc6a1*^em2(S295L)Smoc^, catalog number NM-KI-190014), which performed targeted CRISPR/ Cas9 *Slc6a1* variant generation, the process of which is briefly summarized below. A CC to TA point-mutation was introduced in the *Slc6a1* gene (Gene Ensembl ID: *Slc6a1*-201 ENSMUST00000032454.7) at the exon 9 locus resulting in serine-to-leucine substitution at amino acid 295 (S295L) of GAT-1. First, Cas9 mRNA and gRNA were produced by *in vivo* transcription, and oligonucleotide donor DNA was synthesized. The knock-in locus sequence is as follows; the underlined letter is the guide RNA (gRNA) target site and the blue font is the mutant site:

agcgatgaagatgtcgactctgaccttgatcagttctgaggttctgat cctctgtctgcagGTGTGGCTTGACGCCG CCACCCA<u>GATCTTCTTCT**CC**TACGGGC</u>TGGGCCTGGGGTCCCTGATTGCTCTGGGAAGCTAC AACTCTTTCCACAACAATGTGTACAG gtgcgagggggcgggctttgaggc ttccttcctggccacgcccctaaacatg

The single-stranded donor oligonucleotide ssODN sequence is as follows; the red font is the base after mutation, and the green font is the synonymous mutation to prevent gRNA secondary cleavage:

GTGTGGCTTGACGCCGCCACCCAGATCTTCTTCT**TA**TACGGGCT**C**GGCCTGGGGTCCCTGATTGCTCTGGG AAGCTACAACTCTTTCCACAACAATGTGTACAG

The mixture of Cas9 mRNA, gRNA, and donor DNA was microinjected into fertilized eggs (C57BL/6J). The injected fertilized eggs were transplanted into pseudo-pregnant mother mice, and F0 generation mice were born around 20 days after transplantation. Because the early cleavage rate of the fertilized egg is very fast, the obtained F0 generation mice were chimeras, which might not have the ability of stable inheritance, and needed to be passaged to obtain F1 generation mice that could be stably inherited.

To identify the homologous recombina F0 mice, the following reagents and conditions were used: PCR forward primer 1: 5’-CCAGGAGGAGGAG AGGAACAGAT-3’, PCR reverse primer 2: 5’-CAAGCCAGGCAGGTAGAGCAGAGA-3’, PCR components per reaction (ul),: ddH2O (13.2), GXL PCR buffer (2), 2.5 mM dNTP (2), reverse primer 2 (10pmol/ ul, 0.5), forward primer 2 (10pmol/ul, 0.5), GXL DNA polymerase (TaKaRa, Code No: R050A, 0.5), tail genomic DNA (1), totalling 20 ul. PCR program: step 1: 94 °C, 3 min; step 2: 98 °C, 15 sec; step 3: 57 °C, 15 sec; step 4: 68 °C, 1 min; repeated steps 2–4 for 35 cycles; step 5: 68 °C, 5 min; step 6: 12 °C hold. To verify the mutation using sequencing, the following primer was used: 5’-CCAGGAGGAGGAGGAGGAACAGAT-3’.

Three positive F0 mice were verified by PCR and sequencing. F0 mice were crossed with C57BL/6J mice to generate F1 mice. Six positive F1 mice were verified by PCR and sequencing described above.

Upon receipt of the mice, the GAT-1^S295L/+^ colony was maintained at the Gladstone Institutes by crossing GAT-1^S295L/+^ mice to C57BL/6J mice (ISMR_JAX: 000664). Littermate siblings of both genotypes (GAT-1^+/+^, GAT-1^S295L/+^) of both sexes were used in each experiment. The specific number of mice used in each experiment are indicated in figure legends; sex is encoded by open circles (females) and closed circles (males) of data points in each figure.

### GAT-1^−/−^ mice

GAT-1^−/−^ mice were obtained through a collaboration with Steven Gray at the University of Texas Southwestern Medical Center. The mouse strain, B6.129S1-*Slc6a1*^tm1Lst^/ Mmucd, RRID:MMRRC_000426-UCD, was originally obtained from the Mutant Mouse Resource and Research Center (MMRRC) at University of California at Davis, an NIH-funded strain repository, and was donated to the MMRRC by Henry Lester, Ph.D., California Institute of Technology. Upon receipt of the mice at the Gladstone Institutes, the GAT-1^−/−^ colony was maintained by crossing GAT-1^+/−^ mice to GAT-1^+/−^ mice. Littermate siblings of three genotypes (GAT-1^+/+^, GAT-1^+/−^, GAT-1^−/−^) of both sexes were used in each experiment. The specific number of mice used in each experiment are indicated in figure legends; sex is encoded by open circles (females) and closed circles (males) of data points in each figure.

### Surgical implantation of wireless telemetry devices for chronic physiology recordings

Forty-nine adult mice (23 male, 26 female, 15.8 ± 0.3 weeks, 23.44 ± 0.51 g) underwent surgical implantation of wireless physiology transmitter devices for chronic electrocorticogram (ECoG) and electromyogram (EMG) recordings, as well as for the measurement of body temperature and locomotor activity. We anesthetized mice with vaporized isoflurane (3% induction, 1-2% maintenance, carried by 100% O_2_ at a flow rate of 2 L/min) and placed them in a stereotaxic frame for chronic ECoG implants as previously described (Holden et al., 2021). Implants were commercially purchased wireless telemetry devices (HD-X02, Data Sciences International, St. Paul, MN) which were configured for chronic ECoG and EMG recordings. The implant surgery was performed as previously described (Holden et al., 2021). Briefly, we implanted an ECoG screw in the skull overlying the primary motor cortical region at the following coordinates: 1.0 mm anterior from Bregma, 2.5 mm lateral from the midline (Paxinos and Franklin, 2012). A ground screw was placed overlying the cerebellum (0.5–1 mm posterior to Lambda and 0.5–1 mm lateral to midline). Two EMG electrodes were placed in the deep parasagittal cervical muscles.

The biocompatible housing enclosing the device battery and transmitter was placed subcutaneously overlying the left flank per manufacturer recommendations. The skin was closed over the entire apparatus. For analgesia, topical lidocaine ointment (5%) was applied prior to incision and extended-release buprenorphine (0.05-0.1 mg/kg s.c.) was administered prior to recovery from anesthesia. Mice recovered for 5–7 days after surgery before the start of recordings. To assess physiology throughout the circadian cycle, we housed our animals using a standard light/dark cycle (7 a.m./7 p.m.). Forty-eight-hour baseline recordings were obtained from each mouse, aligned to the start of a light period. Single-housed mice in their home cages were placed over radio frequency receivers that sent signals to an acquisition computer over Ethernet. Simultaneous ECoG, EMG, body temperature, and locomotor activity signals were digitized and continuously recorded from up to eight mice simultaneously using Ponemah software (SCR_017107), and sampled at 500 Hz. Exclusion criteria: Six mice (2 female and 1 male GAT-1^−/−^, 1 male and 1 female GAT-1^S295L/+^, and 1 GAT-1^+/+^ female) died before recordings began. In addition, 5 recordings were excluded due to unacceptable data quality (malpositioned electrode (1), contamination of ECoG by muscle artifact (2), transmitter failure (2)), and 4 mice died or were humanely euthanized before completing the entire protocol (thus, some n’s reported in baseline and nesting panels are higher than in pharmacology panels).

### Pharmacology

Following acquisition of 48h baseline recordings, each mouse was serially challenged with vehicle and drug injections. We administered intraperitoneal injections of vehicle (normal saline 0.1 ml/10 g body weight) every 12h (q12h) at 7 a.m. and 7 p.m. for 2 days, completing a total of 4 doses followed by ≥48 h wash-out. Next, half of the mice received T-type Ca^2+^ antagonist ethosuximide (Sigma E7138) 200 mg/kg i.p. q12h at 7 a.m. and 7 p.m. for 2 days followed by 48 h washout, then GAT-1 antagonist NO-711 (Sigma N142) (10 mg/ kg i.p. q12h at 7 a.m. and 7 p.m. for 2 days). The other half of the mice received NO-711 first then ethosuximide, with a 48 h wash-out between treatments. Each drug was freshly prepared in normal saline for a total volume of 0.1 ml/10 g body weight each injection, and warmed to > 35 °C prior to injection to minimize its influence on core body temperature.

### Detection of spikes and spike-wave discharge events in ECoG

All ECoG features were normalized to baseline ECoG power, defined by root mean square (RMS) amplitude sampled during a spike-free awake interval from each animal, after removal of DC component of signal (0.2 s time constant). Spikes were detected as events exceeding threshold > 7x RMS amplitude. As such, spikes were counted whether they occurred in the context of a spike-wave discharge (SWD) or during inter-ictal intervals or sleep. SWDs were identified with a semi-automated algorithm as follows. The instantaneous frequency of spikes was projected using a smoothing function (exponential kernel, time constant 0.5 s). Candidate SWD episodes were identified as intervals when the smoothed instantaneous frequency function exceeded a threshold of 3.03 ± 0.02 Hz. Threshold was determined individually for each mouse against the gold standard of visually-identified SWDs, using spontaneous seizures wherever possible in genetically susceptible mice, or provoked SWDs during NO-711 treatment in wildtype mice. Onset time, duration, and spikes per episode of candidate SWD were tallied, then episodes containing < 3 spikes or lasting < 0.5 s were excluded as false detections. Post-hoc sampling of false detections revealed these were enriched for electrical artifacts, sleep spindles, and clusters of sharpened elements during slow wave sleep.

### Behavioral Assessment

Forty-eight naive, non-implanted adult mice (13 male and 12 female GAT-1^+/+^, and 12 male and 11 female GAT-1^S295L/+^, 13.21 ± 0.46 weeks) were assessed on a battery of behavioral assessments. Behavioral testing was carried out during the light period between 7 a.m. and 7 p.m. All mice were transferred to the testing room for acclimation at least one hour prior to experiments. Testing boxes were disinfected by MB-10 (chlorine dioxide, 100 ppm) before and after experiments each day. Between experiments, testing boxes were cleaned by 75% ethanol. All mice were tested in all behavioral tests in the same order. All experimenters were blinded to the genotype of individual mice at the time of experiment.

### Elevated Plus Maze

The Elevated Plus Maze box has two open arms and two closed arms. The length of each arm measured 40 cm from the center hub, the side of the square hub measured 6 cm, the height of the walls of two closed arms measured 20 cm, the height of the elevated plus maze from the floor measured 20 cm. Mice were placed in the hub of the maze and allowed to explore for five minutes. The time in each arm and the total distance travelled were monitored by a video camera (SONY HDR-CX675) and analyzed using tracking software (Noldus, SCR_004074).

### Light-dark Box

The light-dark box apparatus was made of an open field box (60 cm by 60 cm by 30 cm) and a dark chamber box (60 cm by 30 cm by 30 cm) with an entry opening (5 cm by 5 cm) insert. The placement of a dark chamber insert divided the apparatus into a light and a dark chamber. Each mouse was placed into the corner of a light chamber and monitored for 5 minutes by a video camera (SONY HDR-CX675) and analyzed using tracking software (Noldus, SCR_004074).

### Open Field

The open field box was made of opaque white acrylic, bottom square sides measured 60 cm and walls measured 30 cm. Each mouse was placed into the corner of an open field box and allowed to explore freely for 10 minutes. Total and minute-by-minute ambulatory distance and percent time in the center of the box of each mouse were monitored by a video camera (SONY HDR-CX675) and analyzed using tracking software (Noldus, SCR_004074).

### Y maze

The Y maze apparatus consisted of three 40 cm long, 10.5 cm wide, and 15 cm high arms made of opaque white acrylic. Each mouse was placed into the hub and allowed to freely explore for 5 minutes monitored by a video camera (SONY HDR-CX675). An entry was defined as the center of the mouse body extending 5 cm into an arm determined by a tracking software (Noldus, SCR_004074). The chronological order of entries into respective arms was determined. Each time the mouse entered all three arms successively (e.g. A-B-C or A-C-B) was considered an alternation set. Percent correct alternations was calculated by dividing the number of sets by the total number of entries minus two (since the first two entries cannot meet criteria for a set). Mice with 10 or fewer total entries were excluded from spontaneous alternation calculations due to insufficient sample size.

### Tube Test for Social Dominance

The tube test for social dominance was performed as previously described by (Arrant et al., 2016; Voskobiynyk et al., 2021). Mice of the same sex, but opposite genotype, were released into opposite ends of a clear plastic tube and allowed to freely interact. Eventually, one mouse will force the other out of the tube. The first mouse with two feet out of the tube was considered to have lost the match. Each mouse was paired with three different opponents of the opposite genotype, and the winning percentage was calculated for each mouse by dividing the number of wins by the total number of matches. The wins were recorded immediately after each match, and all matches were monitored using a video camera (SONY HDR-CX675) for the investigator’s records.

### Three-Chamber Sociability Test

The three-chamber sociability test was adapted from previously described by (Filiano et al., 2013; Voskobiynyk et al., 2021). The three-chamber apparatus consisted of an opaque white acrylic box with a square bottom side measured 60 cm, walls measured 30 cm. Two transparent acrylic insert dividers with 5 cm by 5 cm openings in the center created three chambers (60 cm by 20 cm). Mice were allowed to freely explore a three-chambered testing apparatus for 10 minutes freely entering three chambers prior the introduction of wire cages containing a novel mouse (adult sex-matched C57Bl/6J) or an empty wire cage. Investigation of the novel mouse and an empty cage was monitored for 10 minutes using a video camera (SONY HDR-CX675) and a video tracking software (Noldus, SCR_004074).

### Nesting

Nest-building behavior was assessed according to published methods (Warmus et al., 2014). Briefly, single-housed mice were placed in a fresh cage, and a new square of compacted nesting material (2 × 2 inches) was weighed and placed in the center of each cage. Nestlets were photographed and scored at 0 h and 48 h according to a modified six-point scale (Warmus et al., 2014) modified from (Deacon, 2012): 0, the nestlet is >99% intact; 1, the nestlet is >90% intact; 2, the nestlet is 50–90% intact; 3, the nestlet is 10-50% intact); 4, the nestlet is <10% intact with nest walls higher than the mouse body on <50% of its circumference; and 5, the nestlet is <10% intact with nest walls higher than the mouse body on >50% of its circumference. For scores 1–3, 0.5 was added if there was an identifiable nesting site. For implanted mice, nesting was assessed one week after surgery, concurrent with the 48 h ECoG baseline. For non-implanted mice, nesting was assessed after completion of the behavioral assessment battery.

### Grip strength

Grip strength was assessed with a force transducer (GSM, Ugo Basile No. 47200-327) equipped with a triangular grip bar attachment (adapted from Nevins et al., 1993; Takeshita et al., 2017). Each mouse was suspended over the apparatus until it gripped the bar with both forelimbs, then the examiner pulled back slowly and steadily until the mouse released its grip. This was repeated 5 times, and the force transduced (in grams) was recorded for each trial. The procedure was repeated with a grid attachment, and force assessed with mouse gripping the grid with four limbs. Mean and maximum force were analyzed for each animal in each configuration.

### Statistical analyses

All numerical values are given as means and error bars are standard error of the mean (SEM) unless stated otherwise. Data sets were evaluated for normality, then parametric or non-parametric tests were chosen as appropriate. Data analysis was performed with GraphPad Prism 7/8 (SCR_002798), and Spike2 (SCR_000903). α = 0.05 defined the threshold for statistical significance per convention.

## ACKNOWLEDGEMENTS

We thank Amber Freed and Neil Hackett for insightful discussions that motivated this work. We thank Frances S. Cho for technical assistance with surgeries and Deanna Necula for technical assistance with data visualization. We thank Shreya Kashyap and Erik Roberson for discussion of the behavior assessment design and analysis. We thank Irene Lew and Zanib Naeem for assistance with animal husbandry. We thank Dan Feng of Shanghai Model Organisms for procuring GAT-1^S295L/+^ mice. We thank Henry Lester at the California Institute of Technology for donating GAT-1^S295L/+^ mice to the Mutant Mouse Resource & Research Centers supported by NIH (MMRC). We thank Steven Gray at the University of Texas Southwestern Medical Center for sharing GAT-1^−/−^ mice. We also thank Francoise Chanut, Frances S. Cho, and Deanna Necula for critical feedback on our manuscript.

---

**Supplemental Figure 1.**
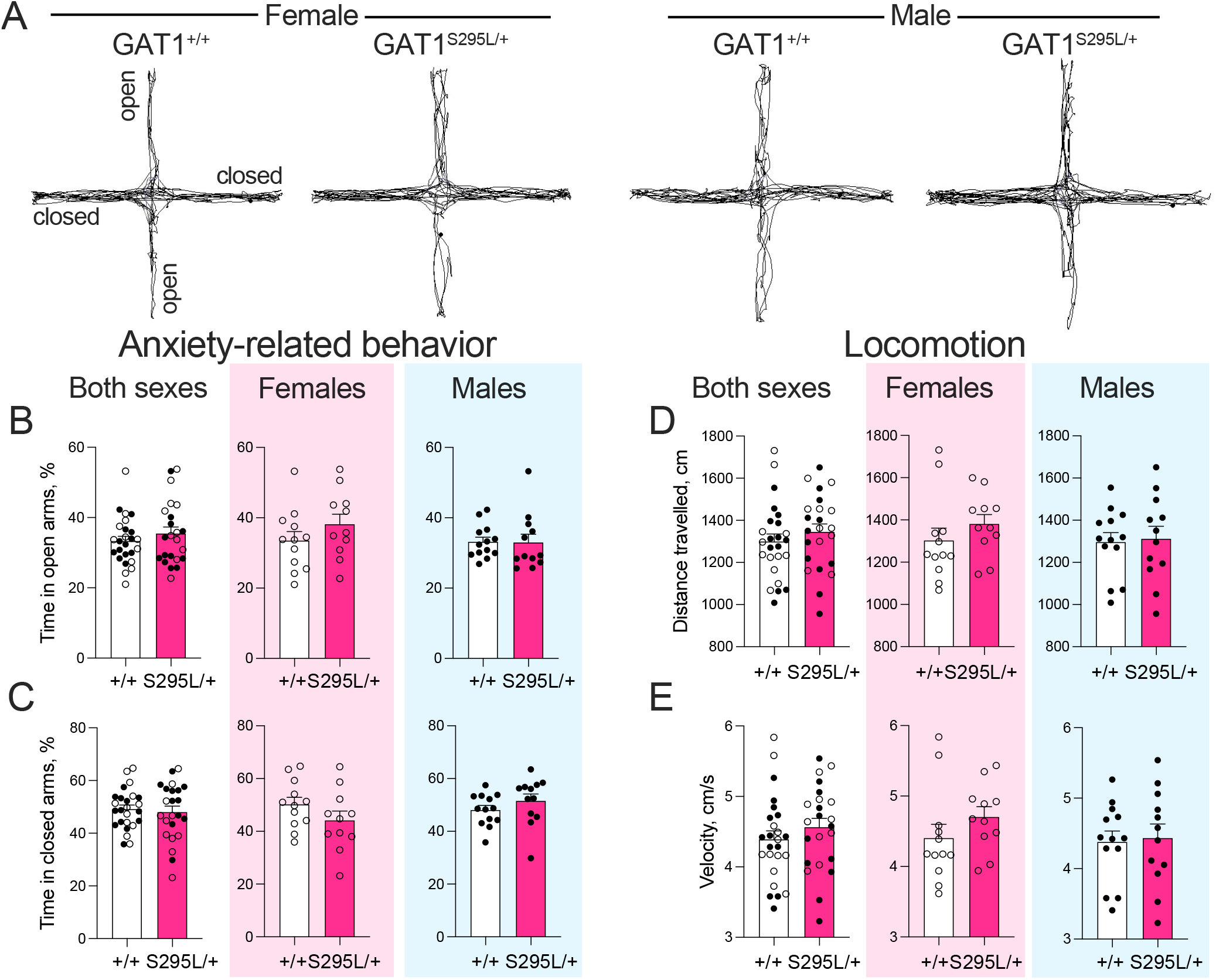
GAT-1^S295L/+^ mice do not exhibit behavioral deficits in the elevated plus maze. **A.** Representative traces of mouse position during 5-min exploration of the elevated plus maze (EPM). **B.** Quantification of proportion of time spent in open arms of EPM (left: unpaired t-test, p=0.3549, n=25,23; middle: unpaired t-test p=0.2364, n=12,11; right: p=0.9510, n=13,12). **C.** Quantification of proportion of time spent in closed arms of EPM (left: unpaired t-test, p=0.6843 n=25,23; middle p=0.1775, n=12,11; right: unpaired t-test, p=0.2677, n=13,12). **D.** Quantification of distance traveled during EPM test (left: unpaired t-test, p=0.3815, n=25,23; middle: unpaired t-test, p=0.3071, n=12,11; right: unpaired t-test, p=0.8280, n=13,12). **E.** Quantification of mean velocity during EPM test. (left: unpaired t-test, p=0.3270, n=25,23; middle: unpaired t-test, p=0.2341, n=12,11; right: unpaired t-test, p=0.8275, n=13,12).

**Supplemental Figure 2.**
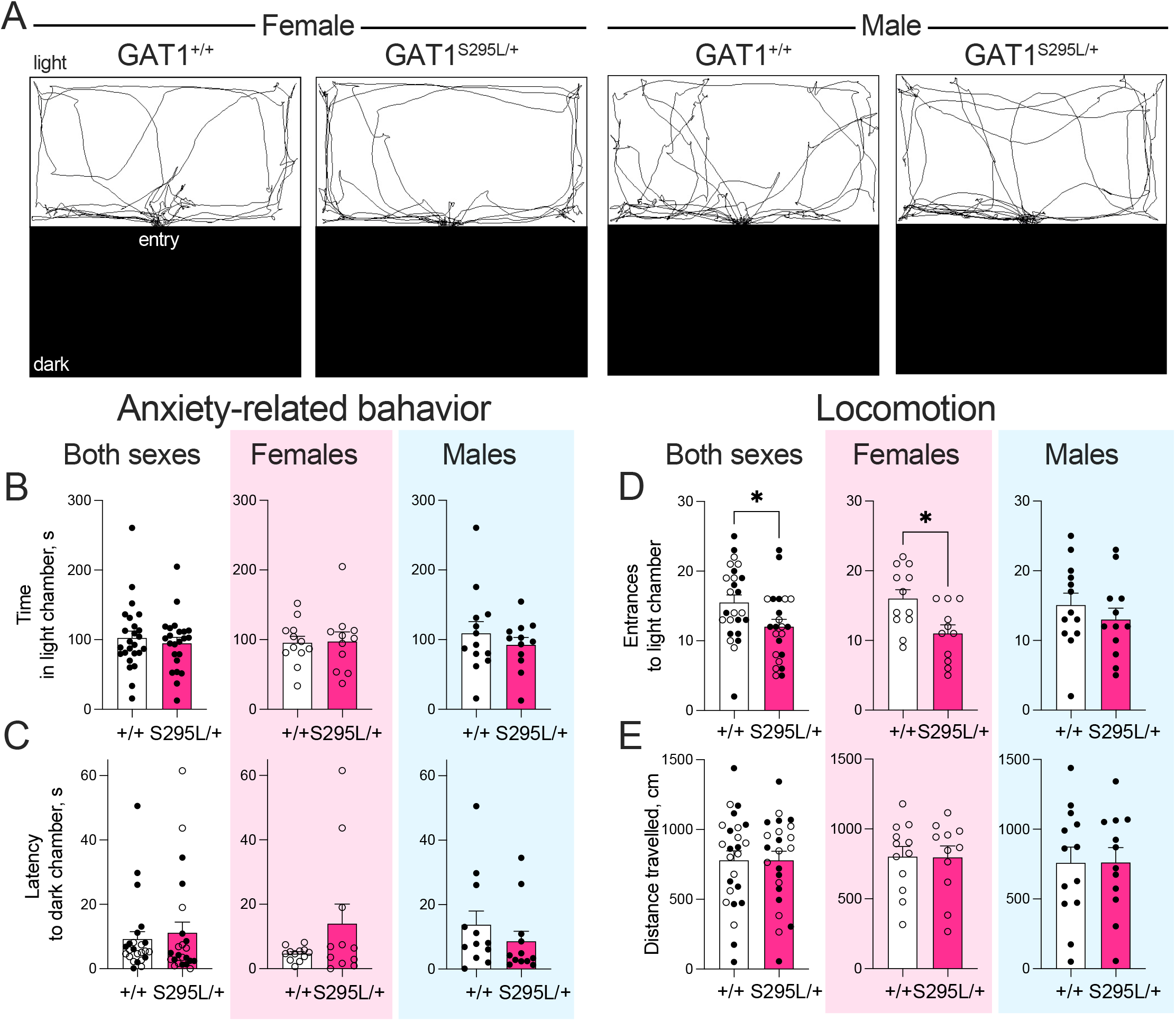
GAT-1^S295L/+^ mice show locomotor deficits, but no anxiety-related behavior, in light-dark box. **A.** Representative tracings of mouse location, tracked over the course of the 5-minute light-dark box assessment. **B.** Quantification of time spent in the light compartment (left: unpaired t-test, p=0.5491, n=25,23; middle: unpaired t-test, p=0.9167, n=12,11; right: unpaired t-test, p=0.4207, n=13,12). **C.** Quantification of latency to move to the dark chamber (left: unpaired t-test, p=0.6229, n=25,23; middle: unpaired t-test, p=0.1251, n=12,11; right: unpaired t-test, p=0.3375, n=13,12). **D.** Quantification of entrances to light chamber (left: unpaired t-test p=0.0235, n=25,23; middle: unpaired t-test, p=0.0105, n=12,11; right: unpaired t-test, p=0.3890, n=13,12). Same as Figure 2A, repeated for clarity. **E.** Quantification of distance traveled during light-dark box test (left: unpaired t-test p=0.9922, n=25,23; middle: unpaired t-test, p=0.9627, n=12,11; right: unpaired t-test, p=0.9843, n=13,12).

**Supplemental Figure 3.**
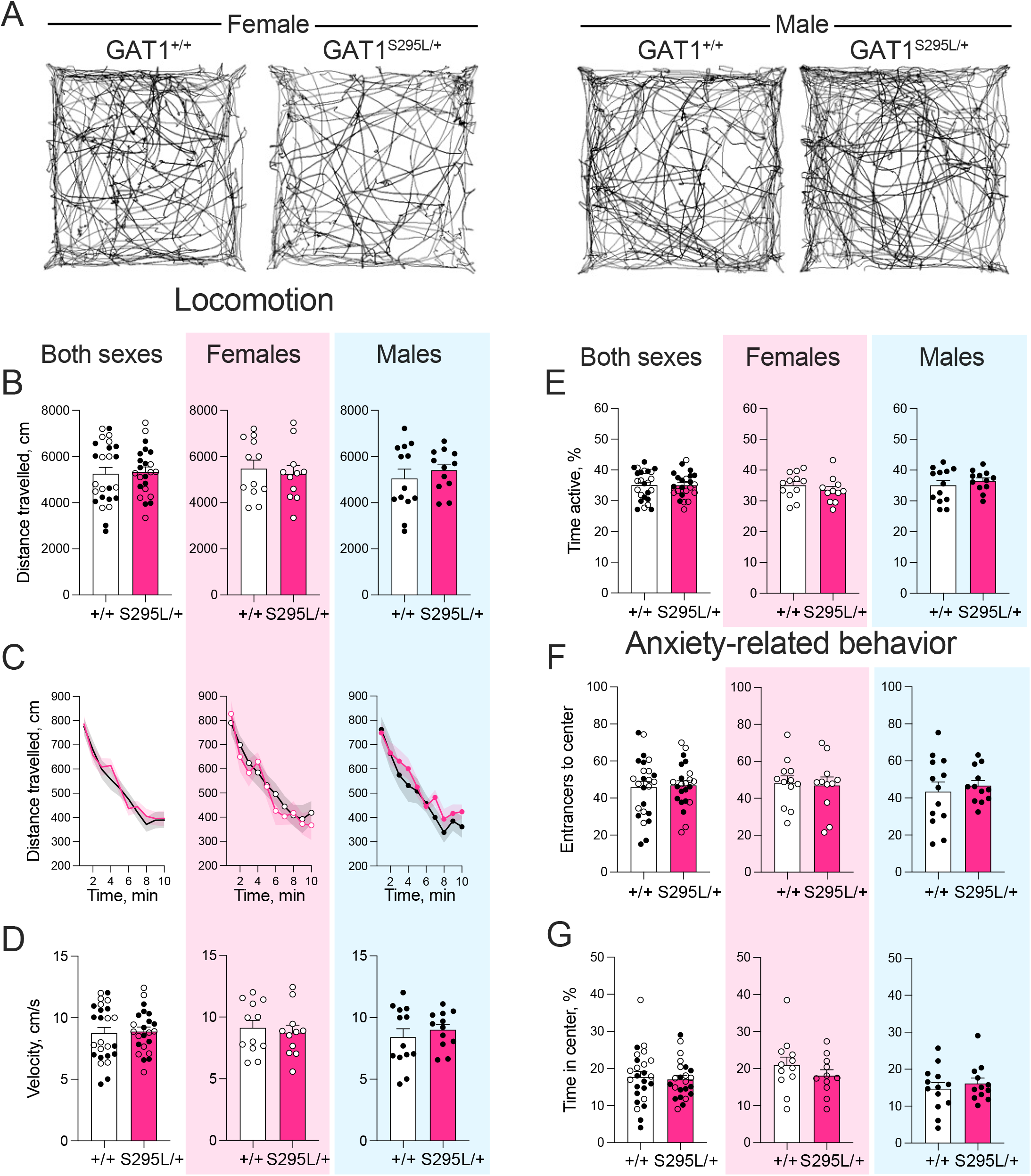
GAT-1^S295L/+^ mice do not exhibit behavioral deficits in the open field during habituation. **A.** Representative plots of mouse location, tracked over the course of the 10-minute open field, performed as the first test in the behavioral battery. **B.** Distance traveled during initial OF testing (left: unpaired t-test p=0.8301, n=25,23; middle: unpaired t-test p=0.6540, n=12,11; right: unpaired t-test p=0.4668, n=13,12). **C.** Quantification of distance traveled as a function of time during initial OF test (left: two-way RM-ANOVA, main effect of genotype p =0.7853, main effect of time p<0.0001; middle: two-way RM-ANOVA, main effect of genotype p 0.7191, main effect of time p<0.0001; right: two-way RM-ANOVA, main effect of genotype p =0.4637, main effect of time p<0.0001). Gray shading indicates the 95% confidence interval. **D.** Mean velocity during initial OF test (left: unpaired t-test p = 0.8302, n=25,23; middle: unpaired t-test p = 0.6539, n=12,11; right: unpaired t-test p = 0.4668, n=13,12). **E.** Proportion of time spent in motion during initial OF test (left: unpaired t-test p =0.9887, n=25,23; middle: unpaired t-test p = 0.3659, n=12,11; right: unpaired t-test p = 0.4394, n=13,12). **F.** Entrances to center during initial OF test (left: unpaired t-test, p=0.8061, n=25,23; middle: unpaired t-test, p=0.8096, n=12,11; right: unpaired t-test, p=0.5912, n=13,12). **G.** Time in center during initial OF test (left: unpaired t-test, p=0.7317, n=25,23; middle: unpaired t-test, p=0.3044, n=12,11; right: unpaired t-test, p=0.5320, n=13,12).

**Supplemental Figure 4.**
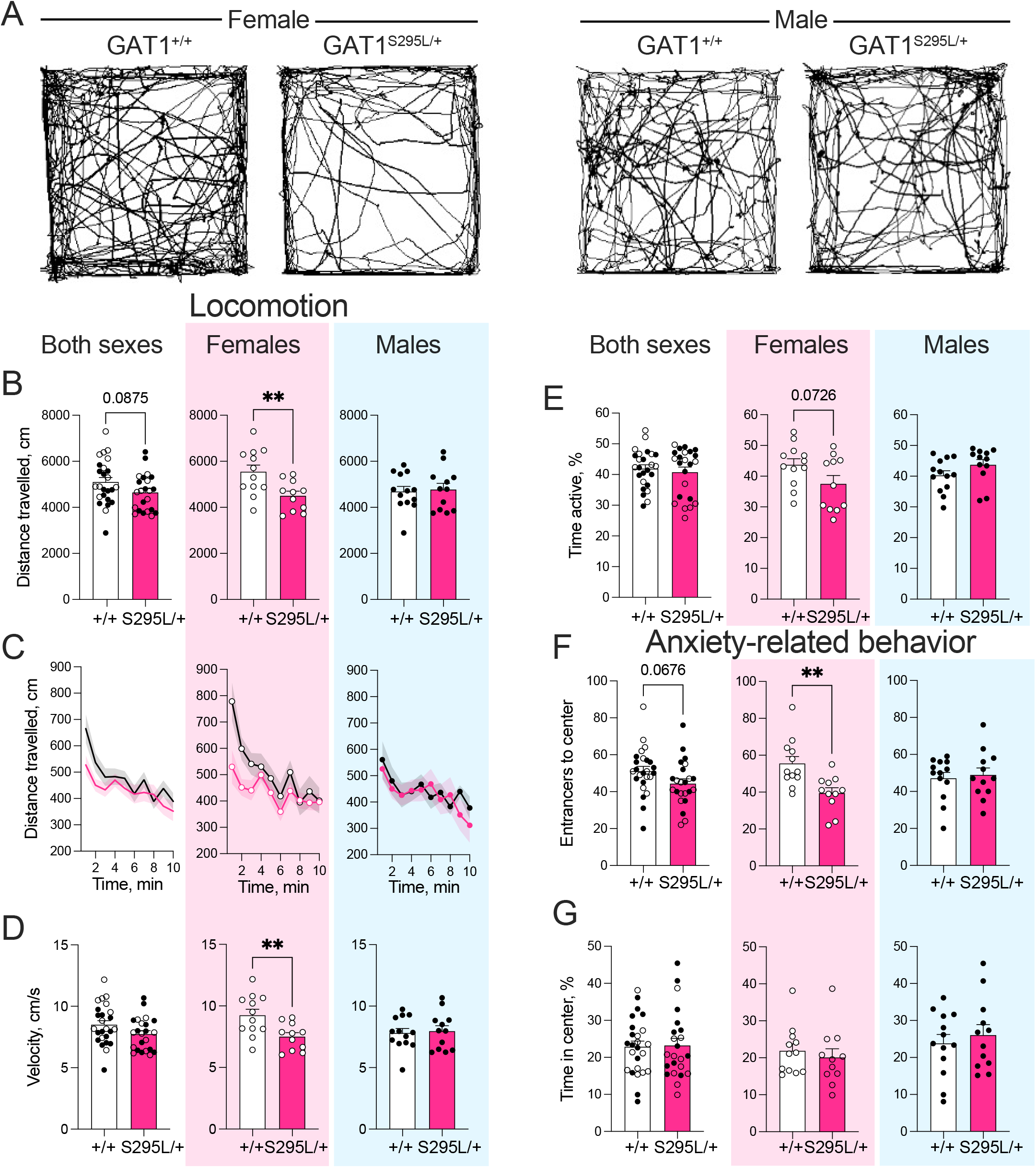
GAT-1^S295L/+^ mice do exhibit behavioral deficits in the open field after habituation. **A.** Representative plots of mouse location, tracked over the course of the 10-minute open field test, repeated at the end of the behavioral battery. Same as Figure 2A, repeated for clarity. **B**. Distance traveled during OF testing (left: unpaired t-test, p=0.0875, n=25,23; middle: unpaired t-test, p = 0.0078, n=12,11; right: unpaired t-test, p = 0.7926, n=13,12). Same as Figure 2B, repeated for clarity. **C.** Quantification of distance traveled as a function of time during repeat OF test (left: twoway RM-ANOVA, main effect of genotype p = 0.1071, main effect of time p < 0.0001, interaction p = 0.2377, n=25,23; middle: two-way RM-ANOVA, main effect of genotype p = 0.0244, main effect of time p < 0.0001, interaction p = 0.0991, n=12,11; right: two-way RM-ANOVA, main effect of genotype p = 0.7420, main effect of time p = 0.055, interaction p = 0.5171, n=13,12). **D.** Quantification of velocity during repeat OF test (left: unpaired t-test, p = 0.0875, n=25,23; middle: unpaired t-test p =0.0078, n=12,11; right: unpaired t-test, p = 0.7922, n=13,12). **E.** Quantification of proportion of time spent in motion during repeat OF test (left: unpaired t-test, p = 0.5739, n=25,23; middle: unpaired t-test, p = 0.0726, n=12,11; right: unpaired t-test, p = 0.1365, n=13,12). **F.** Quantification of entrances to center during repeat OF (left: unpaired t-test, p = 0.0676, n=25,23; middle: unpaired t-test, p = 0.0032, n=12,11; right: unpaired t-test, p = 0.7229, n=13,12). **G.** Quantification of time in center during repeat OF test (left: unpaired t-test, p = 0.8917, n=25,23; middle: unpaired t-test, p = 0.5613, n=12,11; right: unpaired t-test, p = 0.5468, n=13,12).

**Supplemental Figure 5.**
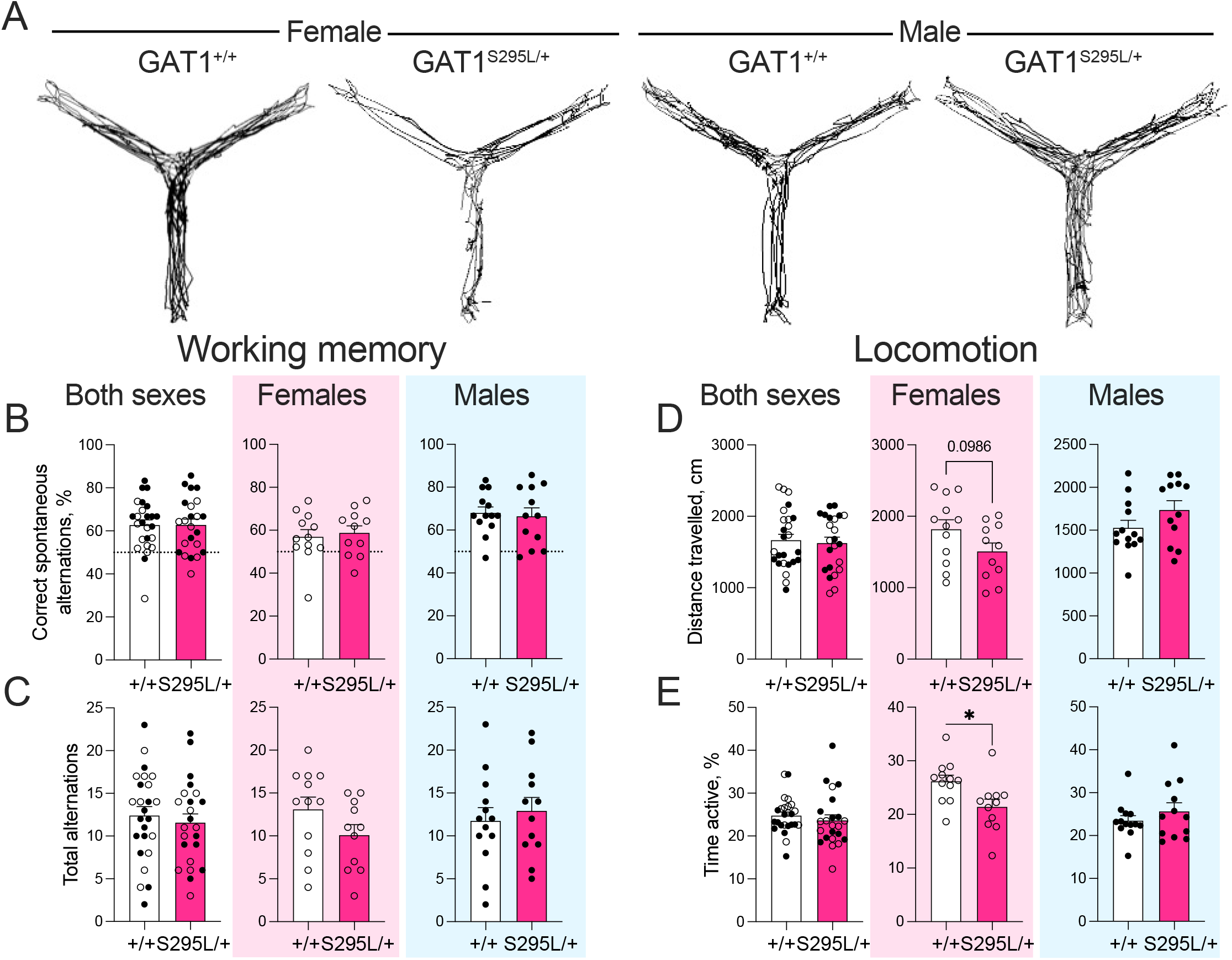
GAT-1^S295L/+^ mice show decreased locomotion, but no working memory deficits in Y maze. **A.** Representative tracings of mouse position over the course of the 5-minute Y-maze test. **B.** Quantification of proportions of spontaneous alterations (left: unpaired t-test, p=0.9779, n=25,23; middle: unpaired t-test, p=0.6905, n=12,11; right: unpaired t-test, p=0.7424, n=13,12). **C.** Quantification of total alternations (left: unpaired t-test, p=0.5727, n=25,23; middle: unpaired t-test, p=0.1322, n=12,11; right: unpaired t-test, p=0.6066, n=13,12). **D.** Quantification of total distance traveled during Y-maze (left: unpaired t-test, p=0.7332, n=25,23; middle: unpaired t-test, p=0.0986, n=12,11; right: unpaired t-test, p=0.1396, n=13,12). **E.** Quantification of proportion of time spent in motion (left: unpaired t-test, p=0.4578, n=25,23; middle: unpaired t-test, p=0.0157, n=12,11; right: unpaired t-test, p=0.3427, n=13,12).

**Supplemental Figure 6.**
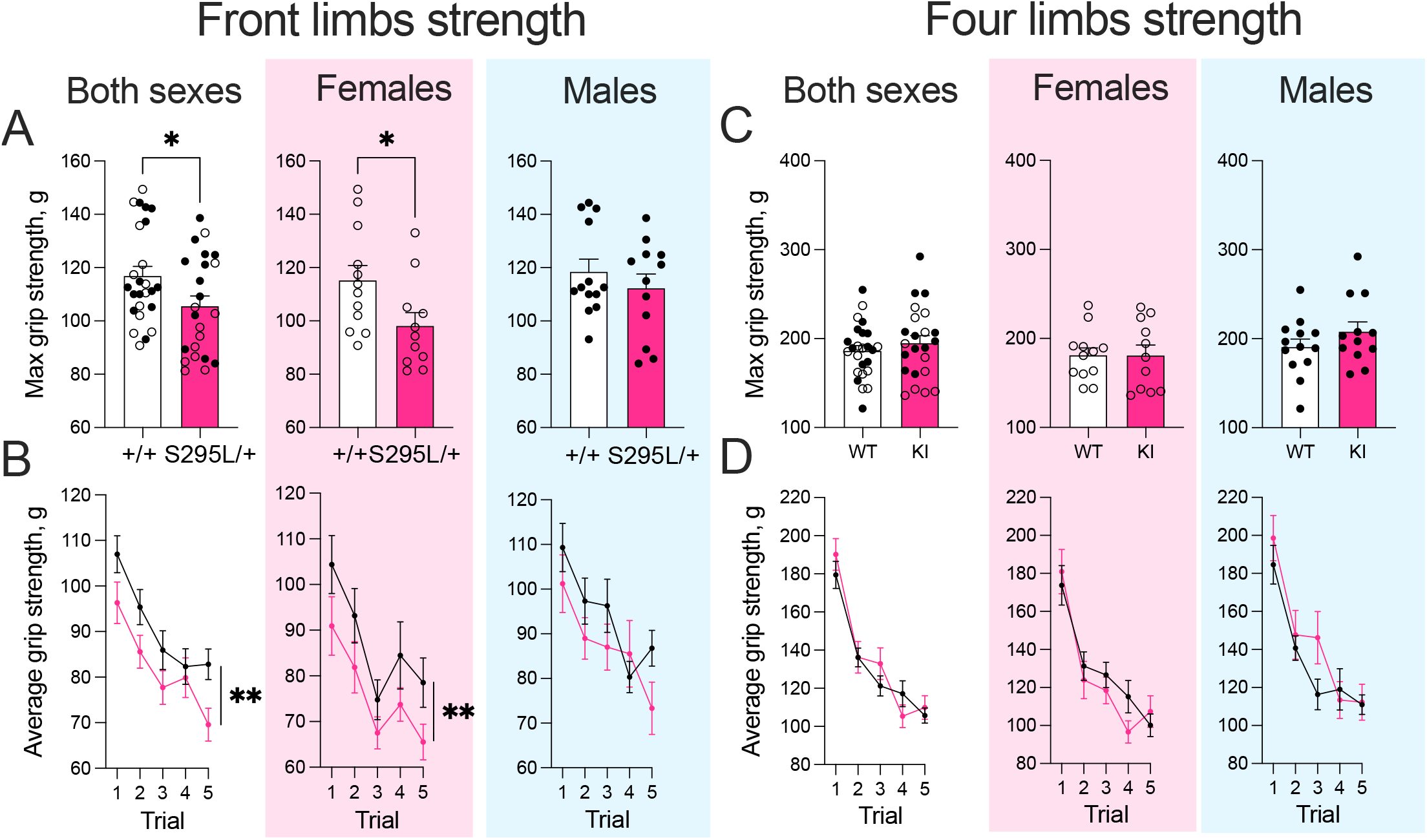
Grip strength is reduced in female but not male GAT-1^S295L/+^ mice. **A.** Quantification of forelimb grip strength, plotted as the best of 5 trials (left: unpaired t-test, n=25,23; middle: unpaired t-test, n=12,11; right: unpaired t-test, p=0.3938, n=13,12). Same as Figure 2A, repeated for clarity. **B.** Quantification of forelimb grip strength over repeated trials (left: two way RM-ANOVA, main effect of genotype p=0.0059, main effect of trial p<0.0001, n=25,23; middle: two way RM-ANOVA, main effect of genotype p=0.0054, main effect of trial p<0.0001, n=12,11; right: two way RM-ANOVA, main effect of genotype p=0.1238, main effect of trial p<0.0001, n=13,12). **C.** Quantification of four limb grip strength, best of 5 trials (left: unpaired t-test, p=0.3918, n=25,23; middle: unpaired t-test, p=0.9961, n=12,11; right: unpaired t-test, p=0.2448, n=13,12). **D.** Quantification of four limb grip strength over repeated trials (left: two way RM-ANOVA, main effect of genotype p=0.6073, main effect of trial p<0.0001, n=25,23; middle: two way RM-ANOVA, main effect of genotype p=0.5585, main effect of trial p<0.0001, n=12,11; right: two way RM-ANOVA, main effect of genotype p=0.3074, main effect of trial p<0.0001, n=13,12).

**Supplemental Figure 7.**
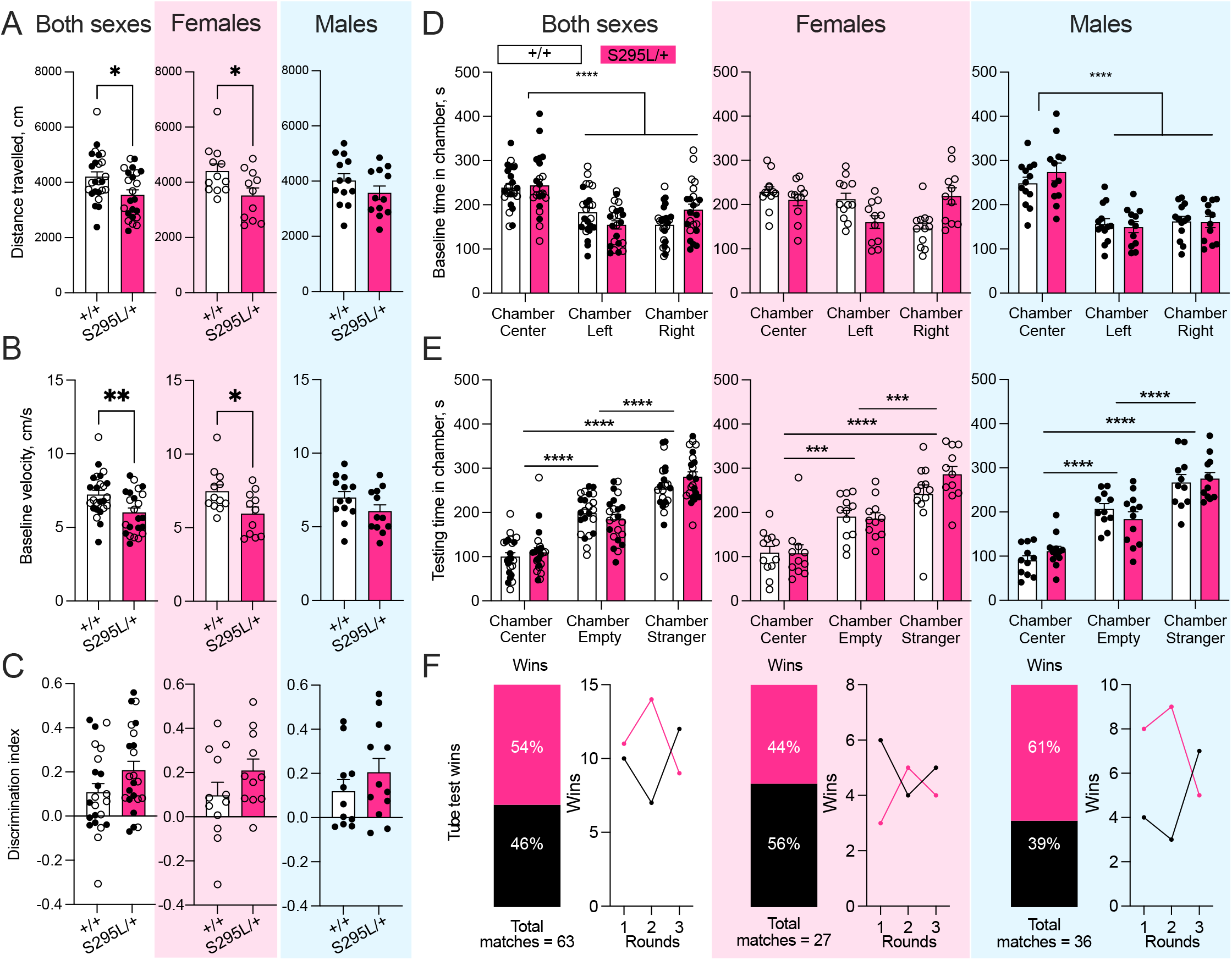
GAT-1^S295L/+^ mice show decreased locomotion, but normal measures of sociability. **A.** Distance travelled in the three-chamber sociability test habituation (left: unpaired t-test, p=0.0118, n=25,23; middle: unpaired t-test, p=0.0258, n=12,11; right: unpaired t-test, p=0.21, n=13,12). Same as Figure 2E, repeated for clarity. **B.** Quantification of velocity in the in the three-chamber sociability test (left: unpaired t-test, p=0.0058, n=25,23; middle: unpaired t-test, p=0.0215, n=12,11; right: unpaired t-test, p=0.1305, n=13,12). **C.** Quantification of discrimination index (proportion of time spent with novel mouse) (left: unpaired t-test, p=0.0809, n=25,23; middle: unpaired t-test, p=0.1701, n=12,11; right: unpaired t-test, p=0.3078, n=13,12). **D.** Time spent in each of three chambers (center, left, and right) during habituation (left: two-way RM-ANOVA, main effect of genotype p=0.2576, main effect of chamber p<0.0001; middle: two-way RM-ANOVA, main effect of genotype p=0.4281, main effect of chamber p=0.0723; right: two-way RM-ANOVA, main effect of genotype p=0.3222, main effect of chamber p<0.0001). **E.** Time spent in each chamber during testing (left: two-way RM-ANOVA, main effect of genotype p=0.2052, main effect of chamber p<0.0001; middle: two-way RM-ANOVA, main effect of genotype p=0.2747, main effect of chamber p<0.0001; right: two-way RM-ANOVA, main effect of genotype p=0.5514, main effect of chamber p<0.0001). F. Tube test wins (left: Binomial test, two-tailed p=0.5323; two-way ANOVA, main effect of genotype p=0.6242, main effect of rounds p>0.9999; middle: Binomial test, two-tailed p=0.7011; two-way ANOVA, main effect of genotype p=0.4778, main effect of rounds p>0.9999; right: Binomial test, two-tailed p=0.2430; two-way ANOVA, main effect of genotype p=0.3828, main effect of rounds p>0.9999).

